# *Clostridium scindens*: an endocrine keystone species in the mammalian gut

**DOI:** 10.1101/2024.08.23.609444

**Authors:** Steven L. Daniel, Jason M. Ridlon

## Abstract

*Clostridium scindens* is a keystone human gut microbial taxonomic group that, while low in abundance, has a disproportionate effect on bile acid and steroid metabolism in the mammalian gut. Numerous studies indicate that the two most studied strains of *C. scindens* (i.e., ATCC 35704 and VPI 12708) are important for a myriad of physiological processes in the host. We focus on both historical and current microbiological and molecular biology work on the Hylemon-Björkhem pathway and the steroid-17,20-desmolase pathway that were first discovered in *C. scindens.* Our most recent analysis now calls into question whether strains currently defined as *C. scindens* represent two separate taxonomic groups. Future directions include developing genetic tools to further explore the physiological role bile acid and steroid metabolism by strains of *C. scindens*, and the causal role of these pathways in host physiology and disease.

## Introduction

Each bacterium that inhabits the human body deserves our attention, and thus its own narrative, and yet we, the perplexed host, must serve, after considerable effort, as the narrator. *Clostridium scindens* has an interesting story and has come into the limelight recently based on renewed interest in secondary bile acids such as deoxycholic acid (DCA) which may play an important role in preventing the vegetative emergence of *Clostridium difficile* in the human gut environment (Buffie et al. 2015, Abt et al. 2016), as well as the role of hydrophobic secondary bile acids in colorectal cancer (O’Keefe 2016, Ocvirk et al. 2021) and hepatocellular carcinoma (Yoshimoto et al. 2013, Ma et al. 2018). Less well known, but likely of equal importance in human physiology and health is the pathway that is the basis for its name “scindens” which means “to cut” owing to the side-chain cleavage of cortisol forming 11-oxy-androgens (Bokkenheuser et al. 1984, Morris et al. 1985, Krafft et al. 1987). *C. scindens* is a core member of the human gut, and perhaps a keystone species, responsible for major biotransformations of bile acids and other steroids that regulate the structure of the gut microbiome and host-microbe interactions. Here, we review the major historical figures and publications relevant to 7ɑ-dehydroxylation of primary bile acids, side-chain cleavage of cortisol, and the isolation and characterization of *C. scindens*, describe the current understanding of steroid metabolism by this bacterial species, host-microbe interactions emerging from this metabolism, and offer some suggestions for future directions.

## Historical paths to *C. scindens*

The path to discovering *C. scindens* began in 1911 with the detection of DCA in human feces by Hans Fischer (Fischer 1911). A series of innovations in chromatography, radiolabeling, and gnotobiology around the mid-20^th^ century confirmed that the removal of the C7-hydroxyl group in vivo was due to microbial action on “primary” bile acids made by the host, which generated “secondary” bile acids (Ridlon et al. 2023).

Two lines of evidence led to the isolation of distinct strains of *C scindens*. The first historical path developed as epidemiological evidence accumulated indicating that populations consuming a “Westernized” diet high in animal protein and fat and low in complex dietary fiber were at an elevated risk for colorectal cancer (CRC) (McGarr et al. 2005). Countries (Sub-Saharan Africa, Japan, and India) consuming a traditional diet high in fiber and resistant starch consistently showed relatively low rates of CRC, as did populations such as Seventh Day Adventists in the United States who consumed a vegetarian diet (McGarr et al. 2005).

By 1970, it was already well established by laboratories in Scandinavia and Japan that fecal bile acids in germ-free animals reflect only those primary bile acid synthesized in the liver, and that bacterial contamination was necessary for detection of secondary bile acids such as DCA and lithocholic acid (LCA) (Ridlon et al. 2023). The work of Bandaru S. Reddy and Ernst Wynder in the 1960s and 1970s identified dietary saturated fat, as compared to oils, which resulted in significant increases in fecal bile acid concentrations (Reddy et al. 1977b). Their pioneering work in rodent models of chemical carcinogenesis established DCA as a tumor-promoter (Reddy et al. 1976, Reddy et al. 1977a). The microbiology of bile acid metabolism lagged during this period, with several reports of successful isolation of bile acid 7α-dehydroxylating bacteria with subsequence loss after transfer or from failure to submit strains to culture collections (Ridlon et al. 2023). In the 1970s, the isolation and study of *Clostridium leptum* strains exhibiting minor biotransformation of cholic acid (CA) to DCA represented the potential to determine the enzymatic basis behind the Samuelsson-Bergstrӧm model (Ridlon et al. 2023). However, bile acid metabolic activity was lost when cell extracts were generated, indicating that separation and purification of “7ɑ-dehydroxylase” and “Δ^6^-reductase” represented, at least under their conditions, a dead end.

In the late 1970s, Rainer Hammann (see **Figure 1** for his photo) at the Institut für Medizinische Microbiologie und Immunologie, Universitat Bonn, Klinkum Venusberg, Bonn, Germany isolated a bacterium from the feces of a colon cancer patient. This strain was sent to the Anaerobic Laboratory at Virginia Polytechnic Institute (VPI) and State University in Blacksburg, Virginia where it was identified by Lillian “Peg” V. Holdeman and W.E.C. “Ed” Moore (**see Figure 1** for their photos) as *Eubacterium* sp. VPI 12708 (Hylemon et al. 1980). In the early 1980s, Holdeman and Moore sent *Eubacterium* sp. VPI 12708 and other strains of gut bacteria to the Hylemon laboratory in the Department of Microbiology and Immunology, Virginia Commonwealth University, Richmond, VA where these strains were screened by Phillip Hylemon, Bryan White (see **Figure 1** for their photos), and others for bile acid metabolism (Phillip Hylemon, personal communication). This research collaboration proved to be quite fruitful as *Eubacterium* sp. VPI 12708 was found to be capable of quantitative conversion of cholic acid (CA) to DCA (Hylemon et al. 1980, White et al. 1980, White et al. 1981, White et al. 1982, White et al. 1983). In these studies, it was possible to characterize bile acid 7α-dehydroxylating activity in both whole cells and CA-induced cell extracts of *Eubacterium* VPI 12708, setting the stage for both the testing of the Samuelsson-Bergström model and the eventual development of the Hylemon-Björkhem Pathway recognized today (Ridlon et al. 2023). *Eubacterium* VPI 12708, twenty years after its initial isolation, would later be reclassified based on 16S rRNA sequencing and DNA–DNA similarity tests as *C. scindens* VPI 12708 by researchers at the Japanese Collection of Microorganisms, RIKEN (Kitahara et al. 2000).

**Figure 1.**
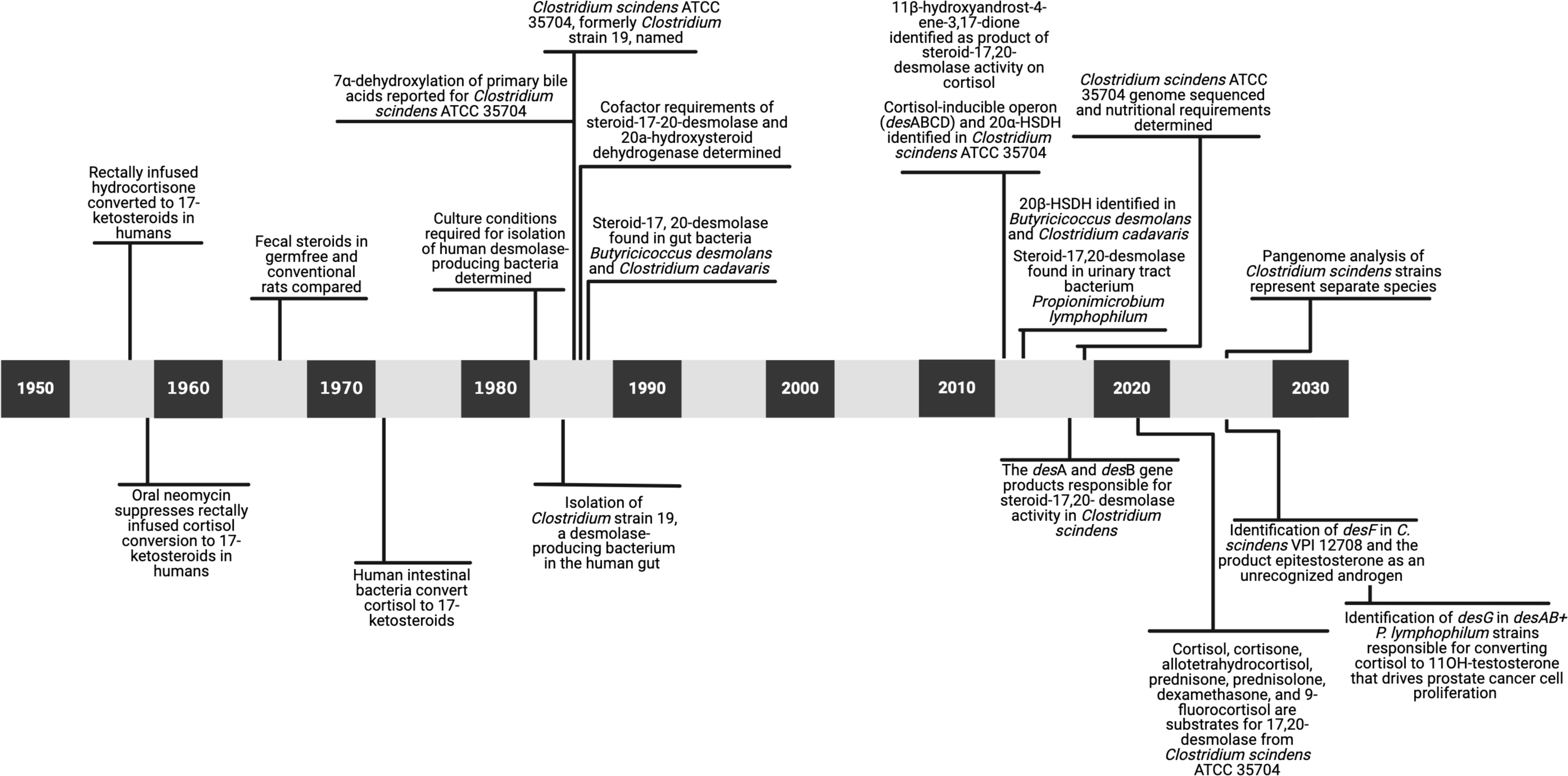
Investigators who worked on the side chain cleavage of steroids, dehydroxylation of bile acids by human fecal bacteria, and isolation and identification of the model gut bacteria *Clostridium scindens* ATCC 35704 and VPI 12708.

A few years after initial reports of *Eubacterium* VPI 12708, a gram-positive spore-forming anaerobe was isolated from the feces of a healthy adult human and named *C. scindens* ATCC 35704^T^ (Bokkenheuser et al. 1984, Morris et al. 1985). This strain, originally designated “*Clostridium* strain 19”, was isolated and characterized by Victor D. Bokkenheuser, Jeanette E. Winter, George N. Morris, Anna M. Cerone-McLernon, Sheryl O’Rourke-Locascio, and others in the Bokkenheuser laboratory in the Department of Pathology, St. Luke’s-Roosevelt Hospital Center, New York, New York, in collaboration with Lillian V. Holdeman and Elizabeth P. Cato at the Anaerobe Laboratory and Alfred E. Ritchie at the National Animal Disease Center in Ames, Iowa (see **Figure 1** for their photos). In contrast to the characterization of *Eubacterium* sp. VPI 12708 due to its ability to convert CA to DCA, *Clostridium* strain 19 was isolated based on selection for its ability to cleave the side-chain of cortisol, forming 11β-hydroxyandrostenedione; it is also capable of bile acid 7α-dehydroxylation of CA (Bokkenheuser et al. 1984, Winter et al. 1984, Morris et al. 1985) (**Figure 2**). This line of research on side-chain cleavage began with reports in the 1950s which indicated that rectal infusions of cortisol in patients with ulcerative colitis resulted in a substantial increase in urinary 17-ketosteroids, which was ablated by oral neomycin treatment (Nabarro et al. 1957, Wade et al. 1959) (**Figure 2**). However, not until 1971 with work by Eriksson and Gustafsson at the Karolinska Institute in Sweden that gas chromatography-mass spectrometry of C-21 corticosteroid incubated with human intestinal contents resulted in confirmation of bacterial side-chain cleavage (Eriksson et al. 1971). In 1981, the Bokkenheuser labidentified bacterial metabolism of cortisol to both C-19 (5β-androstane-3α,11β,17β-triol and 5α-androstane-3α,11β-diol-17-one) and C-21 (tetrahydrocortisol, 21-deoxycortisol, and tetrahydro-21-deoxycortisol) metabolites in human fecal suspensions (Cerone-McLernon et al. 1981) (**Figure 2**). After the report of *Clostridium* strain 19 and description of side-chain cleavage of cortisol, Victor Bokkenheuser collaborated with Phillip Hylemon and Amy Krafft (see **Figure 1** for their photos) and, in a series of papers, a description of the growth and metabolism of cortisol as well as enzymatic activity parameters were described for steroid-17,20-desmolase in *Clostridium* strain 19 (Krafft et al. 1987, Krafft et al. 1989) (**Figure 2**).

**Figure 2.**
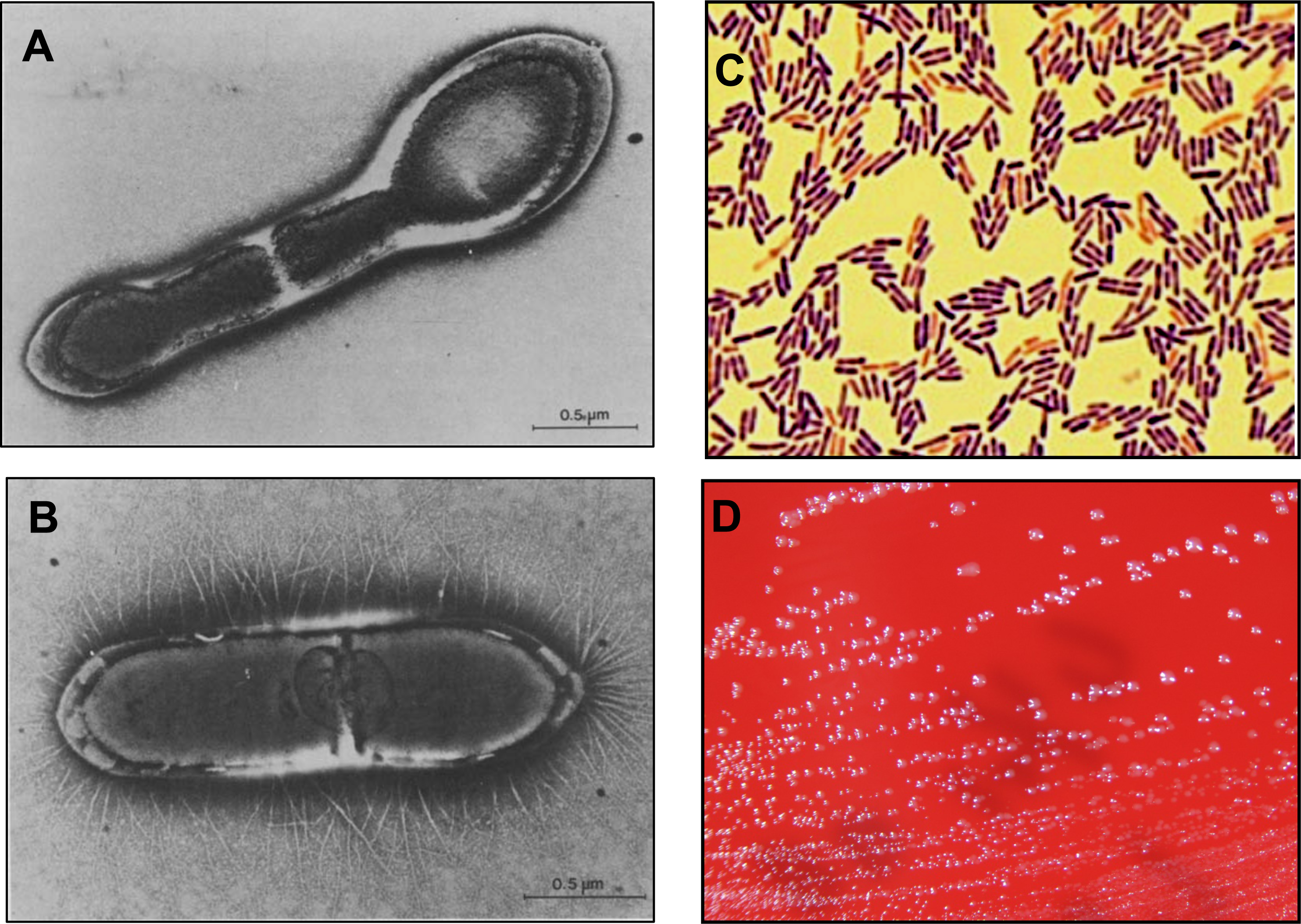
Timeline in the study of bacterial steroid-17,20-desmolase.

So how and when did these historical paths to *C. scindens* merge? In other words, how did *Eubacterium* VPI 12708, an organism capable of bile acid dehydroxylation but not side-chain cleavage, become *C*. *scindens* VPI 12708? In 2000, *Eubacterium* VPI 12708 and 5 additional bile acid-dehydroxylating strains were all reclassified as *C. scindens* based on carbohydrate fermentation profiles, 16S rRNA sequencing (>97% similarity), and DNA–DNA similarity tests. Steroid-metabolizing activities, with the exception of bile acid dehydroxylation (i.e., presence or absence of *bai* genes), were not considered in strain reassignment. A decade earlier, Bokkenheuser and associates who had isolated *C. scindens* ATCC 35704 argued that steroid-metabolizing activities (e.g, 17,20-desmolase activity) are species specific and as such represent distinctive traits that are useful in bacterial identification and taxonomy (Bokkenheuser 1993). Thus, if we were to use these steroid biochemical activities as taxonomy, then organisms capable of bile acid dehydroxylation but not cortisol side-chain cleavage might be more accurately designated in a way to differentiate them from organisms (i.e., *C. scindens*) that both dehydroxylate bile acids and side-chain cleave cortisol. As we will soon demonstrate, we have attempted via pangenome analysis to update and extend the definition of what it might mean to be “*C. scindens”*.

## Taxonomy, morphology, physiology, nutrition, and antibiotic susceptibility of *C. scindens*

*C. scindens* is a mesophilic, chemoheterotrophic, endospore-forming obligately anaerobic bacterium that has been assigned to the following taxa: *Bacillota* (phylum); *Clostridia* (class); *Eubacteriales* (order); *Clostridiaceae* (family); cluster XIVa of *Clostridium* (genus); *scindens* (species); and ATCC 35704; Bokkenheuser 19; CIP 106687; DSM 5676; JCM 6567 (type strain) (Collins et al. 1994, Kitahara et al. 2000, Parte et al. 2020). Cells of *C. scindens* ATCC 35704 are nonmotile, non-flagellated, often fimbriated, occur as Gram-positive rods singly or in chains, and form terminal spores (**Figure 3**) (Bokkenheuser et al. 1984, Morris et al. 1985). Relative to colonial morphology, colonies on blood agar plates are nonhemolytic, convex, smooth, glistening, and white with an entire margin (**Figure 3**).

**Figure 3.**
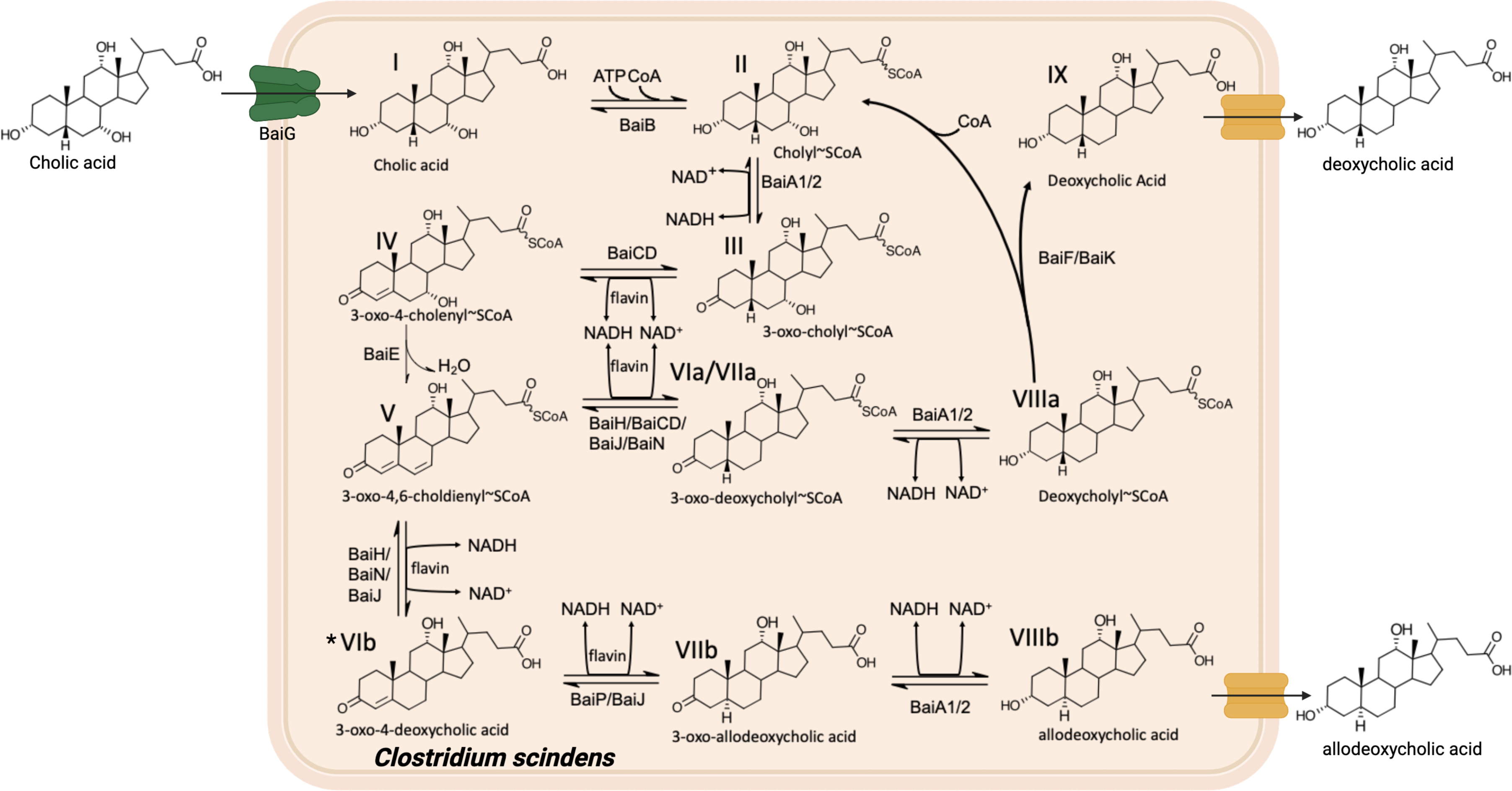
Colony and cellular morphology of *Clostridium scindens* ATCC 35704. (A and B) Electron micrographs of *Clostridium scindens* ATCC 35704 (Bokkenheuser et al. 1984). Used with kind permission from Oxford University Press. Gram stain (C) of cells and colonies (D) of *Clostridium scindens* ATCC 35704 grown on anaerobic EG agar after three days of incubation. Used with kind permission from RIKEN and the Japan Collection of Microorganisms.

As a saccharolytic bacterium, *C. scindens* ATCC 35704 utilizes 6-carbon monosaccharides (glucose, fructose, mannose, and galactose), 5-carbon monosaccharides (ribose and xylose), 6-carbon sugar alcohols (dulcitol and sorbitol), and a disaccharide (lactose) for fermentation and growth (Bokkenheuser et al. 1984, Morris et al. 1985, Kitahara et al. 2000, Devendran et al. 2019). Glucose fermentation proceeds via the Embden-Meyerhof-Parnas (EMP) pathway and typically yields ethanol, acetate, formate, and H_2_ gas as major end products (>1 mM) and succinate, lactate, isobutyrate, and isovalerate as minor end products (<1 mM) (Morris et al. 1985, Devendran et al. 2019). *C. scindens* does not produce lecithinase, lipase, or catalase and is unable to digest gelatin, milk, or meat. *C. scindens* ATCC 35704 is incapable of nitrate reduction, or hydrolysis of starch or esculin. Hydrogen sulfide is produced in sulfide-indole motility medium. See **Table 1** for more information on the overall metabolic profile of *C*. *scindens* ATCC 35704 as well as other strains of *C. scindens* including VPI 12708.

**Table 1.**
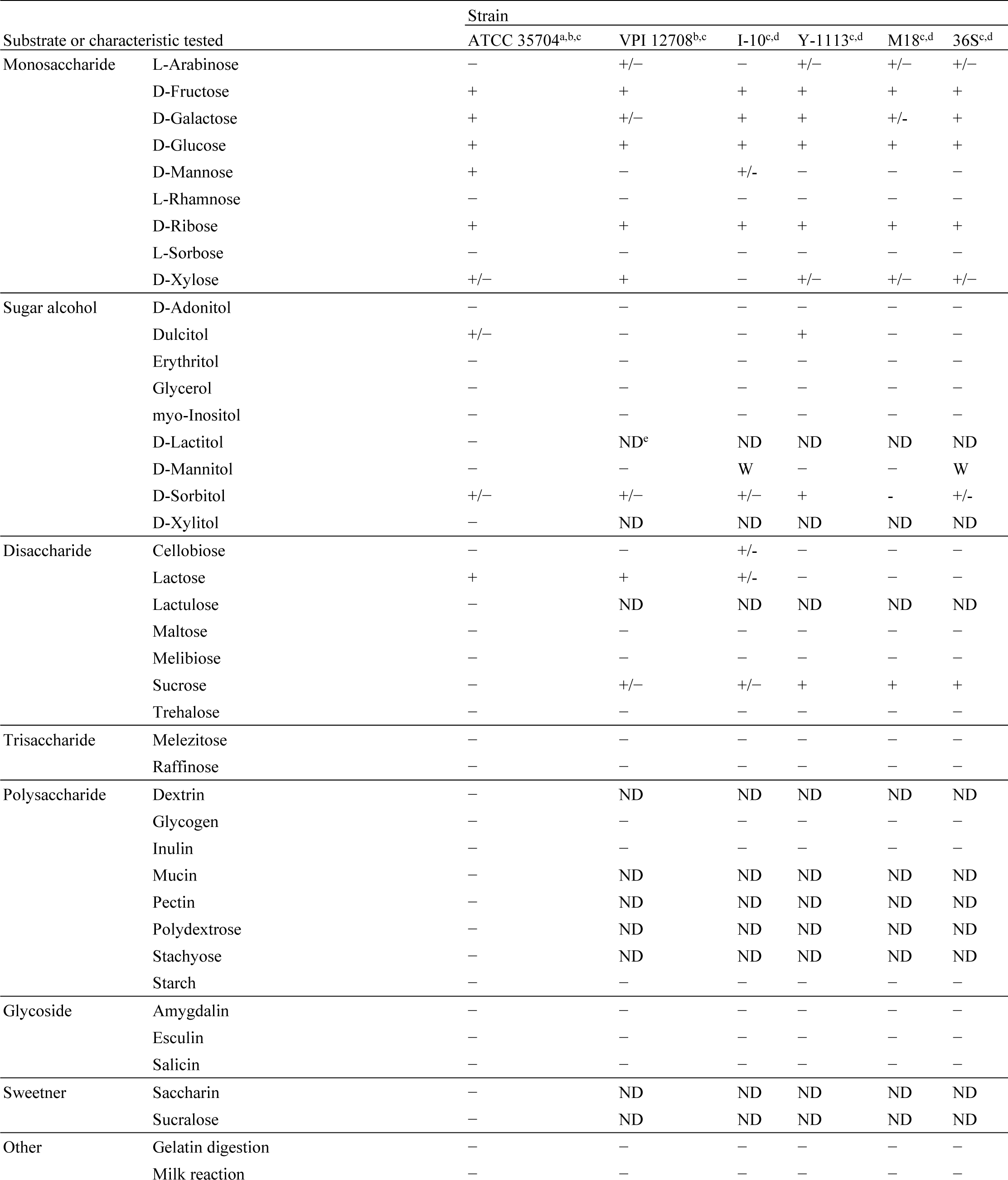
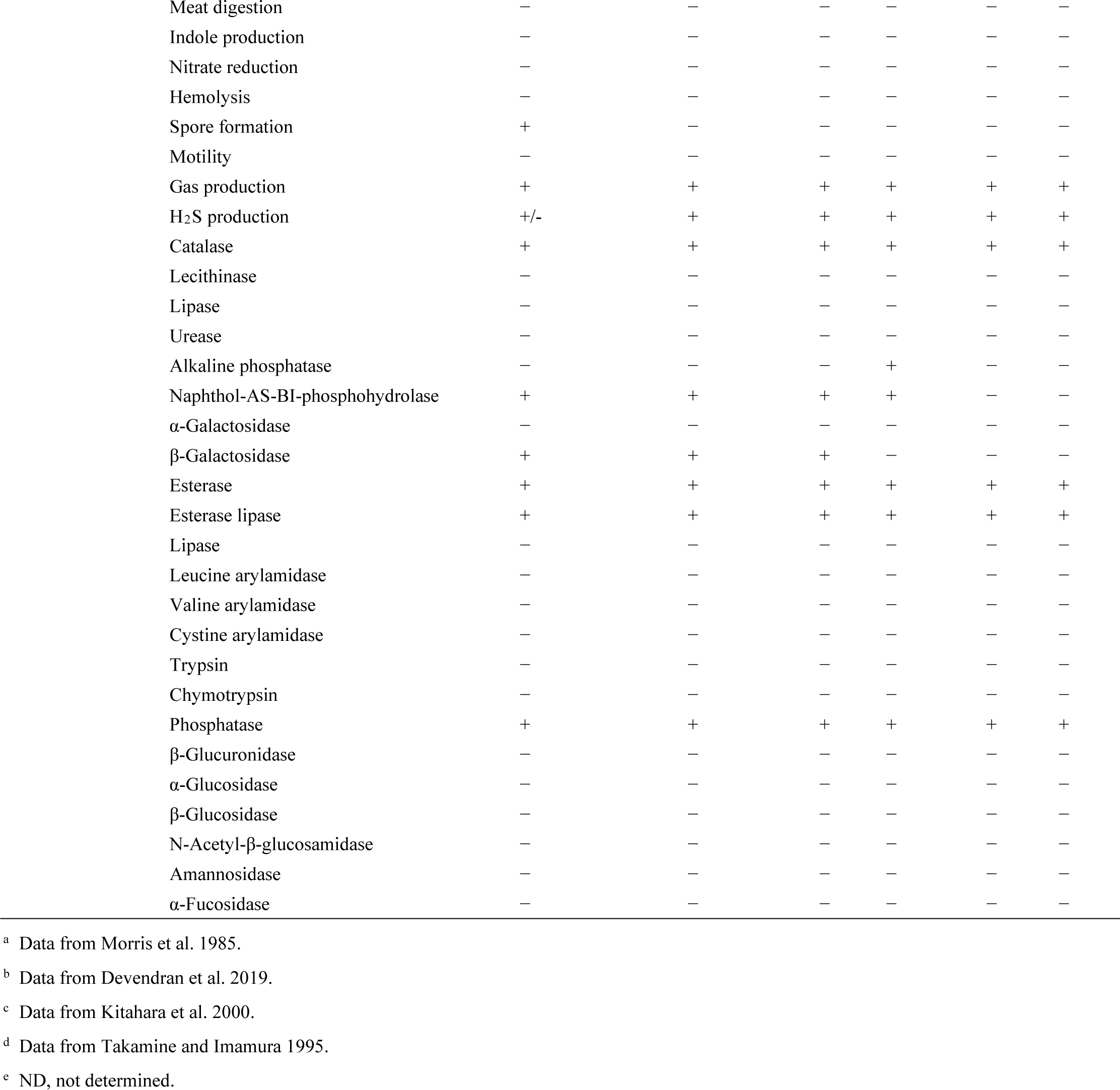
Metabolic profiles of *Clostridium scindens* strains.

| Substrate or characteristic tested |  | Strain |  |  |  |  |  |
| --- | --- | --- | --- | --- | --- | --- | --- |
|  |  | ATCC 35704<sup>a,b,c</sup> | VPI 12708<sup>b,c</sup> | I-10<sup>c,d</sup> | Y-1113<sup>c,d</sup> | M18<sup>c,d</sup> | 36S<sup>c,d</sup> |
| Monosaccharide | L-Arabinose | – | +/- | – | +/- | +/- | +/- |
|  | D-Fructose | + | + | + | + | + | + |
|  | D-Galactose | + | +/- | + | + | +/- | + |
|  | D-Glucose | + | + | + | + | + | + |
|  | D-Mannose | + | – | +/- | – | – | – |
|  | L-Rhamnose | – | – | – | – | – | – |
|  | D-Ribose | + | + | + | + | + | + |
|  | L-Sorbose | – | – | – | – | – | – |
|  | D-Xylose | +/- | + | – | +/- | +/- | +/- |
| Sugar alcohol | D-Adonitol | – | – | – | – | – | – |
|  | Dulcitol | +/- | – | – | + | – | – |
|  | Erythritol | – | – | – | – | – | – |
|  | Glycerol | – | – | – | – | – | – |
|  | myo-Inositol | – | – | – | – | – | – |
|  | D-Lactitol | – | ND<sup>e</sup> | ND | ND | ND | ND |
|  | D-Mannitol | – | – | W | – | – | W |
|  | D-Sorbitol | +/- | +/- | +/- | + | – | +/- |
|  | D-Xylitol | – | ND | ND | ND | ND | ND |
| Disaccharide | Cellobiose | – | – | +/- | – | – | – |
|  | Lactose | + | + | +/- | – | – | – |
|  | Lactulose | – | ND | ND | ND | ND | ND |
|  | Maltose | – | – | – | – | – | – |
|  | Melibiose | – | – | – | – | – | – |
|  | Sucrose | – | +/- | +/- | + | + | + |
|  | Trehalose | – | – | – | – | – | – |
| Trisaccharide | Melezitose | – | – | – | – | – | – |
|  | Raffinose | – | – | – | – | – | – |
| Polysaccharide | Dextrin | – | ND | ND | ND | ND | ND |
|  | Glycogen | – | – | – | – | – | – |
|  | Inulin | – | – | – | – | – | – |
|  | Mucin | – | ND | ND | ND | ND | ND |
|  | Pectin | – | ND | ND | ND | ND | ND |
|  | Polydextrose | – | ND | ND | ND | ND | ND |
|  | Stachyose | – | ND | ND | ND | ND | ND |
|  | Starch | – | – | – | – | – | – |
| Glycoside | Amygdalin | – | – | – | – | – | – |
|  | Esculin | – | – | – | – | – | – |
|  | Salicin | – | – | – | – | – | – |
| Sweetner | Saccharin | – | ND | ND | ND | ND | ND |
|  | Sucralose | – | ND | ND | ND | ND | ND |
| Other | Gelatin digestion | – | – | – | – | – | – |
|  | Milk reaction | – | – | – | – | – | – |

|  |  |  |  |  |  |  |
| --- | --- | --- | --- | --- | --- | --- |
| Meat digestion | – | – | – | – | – | – |
| Indole production | – | – | – | – | – | – |
| Nitrate reduction | – | – | – | – | – | – |
| Hemolysis | – | – | – | – | – | – |
| Spore formation | + | – | – | – | – | – |
| Motility | – | – | – | – | – | – |
| Gas production | + | + | + | + | + | + |
| H<sub>2</sub>S production | +/- | + | + | + | + | + |
| Catalase | + | + | + | + | + | + |
| Lecithinase | – | – | – | – | – | – |
| Lipase | – | – | – | – | – | – |
| Urease | – | – | – | – | – | – |
| Alkaline phosphatase | – | – | – | + | – | – |
| Naphthol-AS-BI-phosphohydrolase | + | + | + | + | – | – |
| α-Galactosidase | – | – | – | – | – | – |
| β-Galactosidase | + | + | + | – | – | – |
| Esterase | + | + | + | + | + | + |
| Esterase lipase | + | + | + | + | + | + |
| Lipase | – | – | – | – | – | – |
| Leucine arylamidase | – | – | – | – | – | – |
| Valine arylamidase | – | – | – | – | – | – |
| Cystine arylamidase | – | – | – | – | – | – |
| Trypsin | – | – | – | – | – | – |
| Chymotrypsin | – | – | – | – | – | – |
| Phosphatase | + | + | + | + | + | + |
| β-Glucuronidase | – | – | – | – | – | – |
| α-Glucosidase | – | – | – | – | – | – |
| β-Glucosidase | – | – | – | – | – | – |
| N-Acetyl-β-glucosamidase | – | – | – | – | – | – |
| Amannosidase | – | – | – | – | – | – |
| α-Fucosidase | – | – | – | – | – | – |

<sup>a</sup> Data from Morris et al. 1985.
<sup>b</sup> Data from Devendran et al. 2019.
<sup>c</sup> Data from Kitahara et al. 2000.
<sup>d</sup> Data from Takamine and Imamura 1995.
<sup>e</sup> ND, not determined.

*C. scindens* ATCC 35704 has by tradition been cultivated under strict anaerobic conditions at 37°C and a pH between 6.5 and 7 in highly enriched, culture media (e.g., chopped meat medium, supplemented brain heart infusion broth, or supplemented peptone broth). Efforts have been made recently to define the nutritional requirements of *C. scindens* ATCC 35704 (Devendran et al. 2019). This initially required adapting *C. scindens* ATCC 35704 to a CO_2_–bicarbonate buffered defined medium (DM) that contained minerals, glucose, vitamins, and amino acids (**Table 2**). Once adapted to DM, the leave-one-amino-acid-group-out and leave-one-vitamin-out approaches were used to resolve the vitamin and amino acid requirements for *C. scindens* ATCC 35704. Riboflavin, pantothenic acid, and pyridoxal•HCl are the sole vitamins and tryptophan is the sole amino acid required for growth by *C. scindens* ATCC 35704 (Devendran et al. 2019). Indeed, genomic analysis supports these findings since genes for tryptophan, riboflavin, pyridoxal phosphate, and pantothenic acid biosynthesis are absent. A defined medium for *C. scindens* ATCC 35704 provides a valuable tool for the assessment of growth, carbon and reductant flow during carbohydrate fermentation, and steroid metabolism and for the development of a much-needed genetic system in this organism.

**Table 2.**
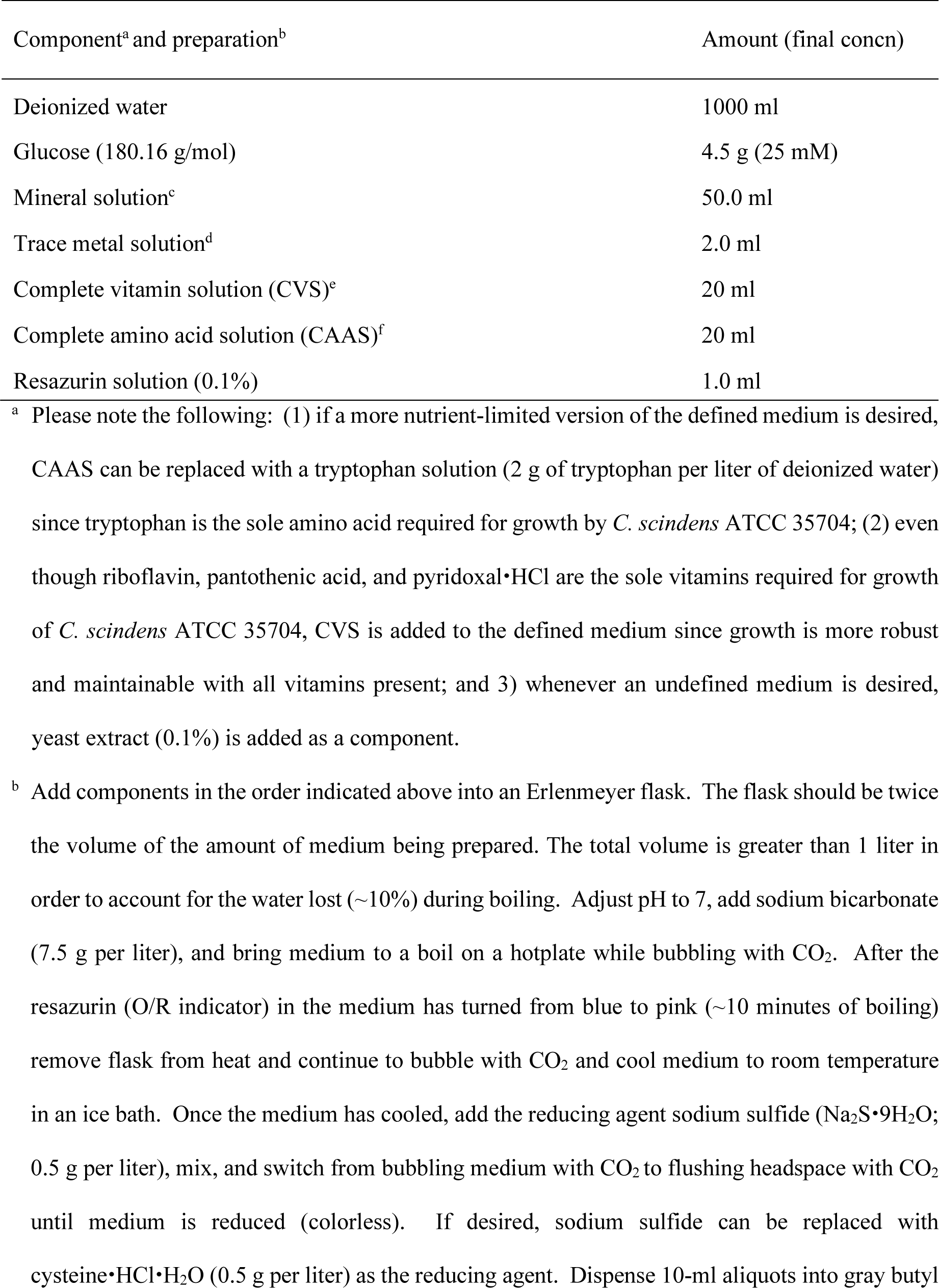
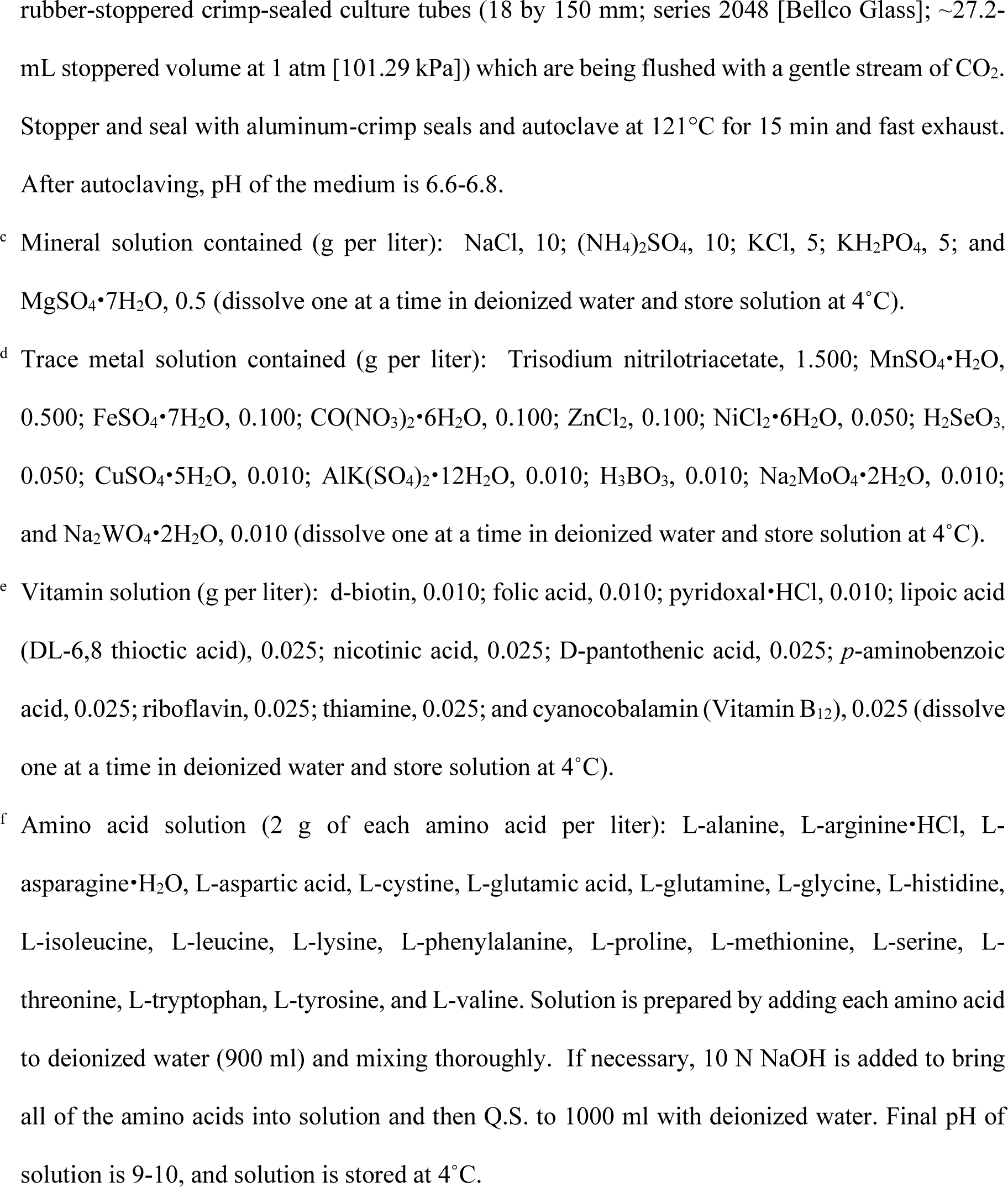
A defined medium for the cultivation of *Clostridium scindens* ATCC 35704.

| Component<sup>a</sup> and preparation<sup>b</sup> | Amount (final concn) |
| --- | --- |
| Deionized water | 1000 ml |
| Glucose (180.16 g/mol) | 4.5 g (25 mM) |
| Mineral solution<sup>c</sup> | 50.0 ml |
| Trace metal solution<sup>d</sup> | 2.0 ml |
| Complete vitamin solution (CVS)<sup>e</sup> | 20 ml |
| Complete amino acid solution (CAAS)<sup>f</sup> | 20 ml |
| Resazurin solution (0.1%) | 1.0 ml |

<sup>a</sup> Please note the following: (1) if a more nutrient-limited version of the defined medium is desired, CAAS can be replaced with a tryptophan solution (2 g of tryptophan per liter of deionized water) since tryptophan is the sole amino acid required for growth by *C. scindens* ATCC 35704; (2) even though riboflavin, pantothenic acid, and pyridoxal•HCl are the sole vitamins required for growth of *C. scindens* ATCC 35704, CVS is added to the defined medium since growth is more robust and maintainable with all vitamins present; and 3) whenever an undefined medium is desired, yeast extract (0.1%) is added as a component.
<sup>b</sup> Add components in the order indicated above into an Erlenmeyer flask. The flask should be twice the volume of the amount of medium being prepared. The total volume is greater than 1 liter in order to account for the water lost (~10%) during boiling. Adjust pH to 7, add sodium bicarbonate (7.5 g per liter), and bring medium to a boil on a hotplate while bubbling with CO<sub>2</sub>. After the resazurin (O/R indicator) in the medium has turned from blue to pink (~10 minutes of boiling) remove flask from heat and continue to bubble with CO<sub>2</sub> and cool medium to room temperature in an ice bath. Once the medium has cooled, add the reducing agent sodium sulfide (Na<sub>2</sub>S•9H<sub>2</sub>O; 0.5 g per liter), mix, and switch from bubbling medium with CO<sub>2</sub> to flushing headspace with CO<sub>2</sub> until medium is reduced (colorless). If desired, sodium sulfide can be replaced with cysteine•HCl•H<sub>2</sub>O (0.5 g per liter) as the reducing agent. Dispense 10-ml aliquots into gray butyl
rubber-stoppered crimp-sealed culture tubes (18 by 150 mm; series 2048 [Bellco Glass]; ~27.2-mL stoppered volume at 1 atm [101.29 kPa]) which are being flushed with a gentle stream of CO<sub>2</sub>. Stopper and seal with aluminum-crimp seals and autoclave at 121°C for 15 min and fast exhaust. After autoclaving, pH of the medium is 6.6-6.8.

Another area which has received little study is the response of *C. scindens* to antimicrobial agents. Using an anaerobic broth-disk method, Morris et al. 1985 reported that *C. scindens* ATCC 35704 was susceptible to penicillin G but resistant to such commonly used antibiotics as tetracycline, chloramphenicol, clindamycin, and erythromycin. Whether other strains of *C. scindens* have similar resistance profiles is unknown. However, it is tempting to speculate that, if resistance among commensal strains of *C. scindens* mirrors that of *C. scindens* ATCC 35704, *C. scindens* would have a competitive advantage during host antimicrobial therapy, thereby allowing it to survive and engage in “ecological suppression” (Waldetoft et al. 2023) of pathogens.

## The bile acid inducible (*bai*) regulon and hydroxysteroid dehydrogenases (HSDHs)

In the late 1950s and early 1960s, the Nobel laureates, Sune K. Bergström and Bengt Samuelsson, performed CA isotope labeling studies in rodents and proposed a diaxial trans-elimination of the 7ɑ-hydroxyl group and 6β-hydrogen followed by reduction of the resultant Δ^6^-intermediate (Ridlon et al. 2023). An important experiment was performed in 1981, a year after the initial reports of bile acid metabolism by *Eubacterium* VPI 12708, that identified multiple CA-inducible polypeptides by one and two-dimensional SDS-PAGE: one at 77 kDa, two at 56 kDa, 27 KDa, and 23.5 KDa (White et al. 1981). Work over the next four decades has resulted in a current model for bile acid 7ɑ-dehydroxylation (**Figure 4**) and gene organization for both bile acid and steroid metabolism by *Clostridium scindens* VPI 12708 and ATCC 35704 strains (**Figure 5**). We have reviewed this history in detail recently and proposed this pathway be named the Hylemon-Bjorkhem Pathway (Ridlon et al. 2023).

**Figure 4.**
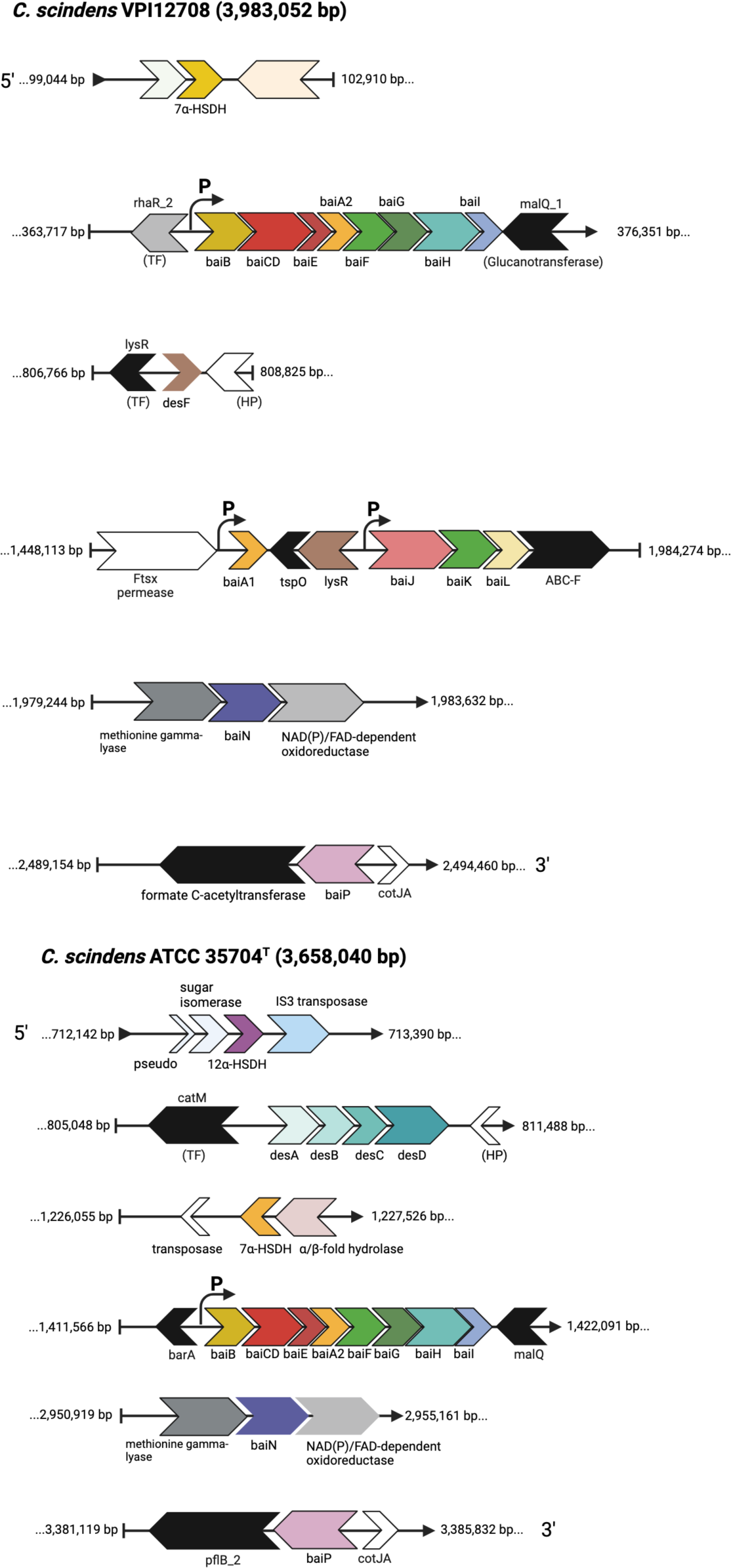
The current model of the Hylemon-Björkhem Pathway of bile acid 7a-dehydroxylation of cholic acid by *Clostridium scindens* strains. Steps V-VIA(VIb) have been shown to be catalyzed by BaiH/BaiN and BaiJ enzymes (). *Step VIb – VIIb has been shown to be catalyzed by BaiJ and BaiP (). Bile acid exporters (orange) have not yet been identified. Each enzymatic step is described in detail in the associated text. Modified from (Devendran et al. 2019) with kind permission.

**Figure 5.**
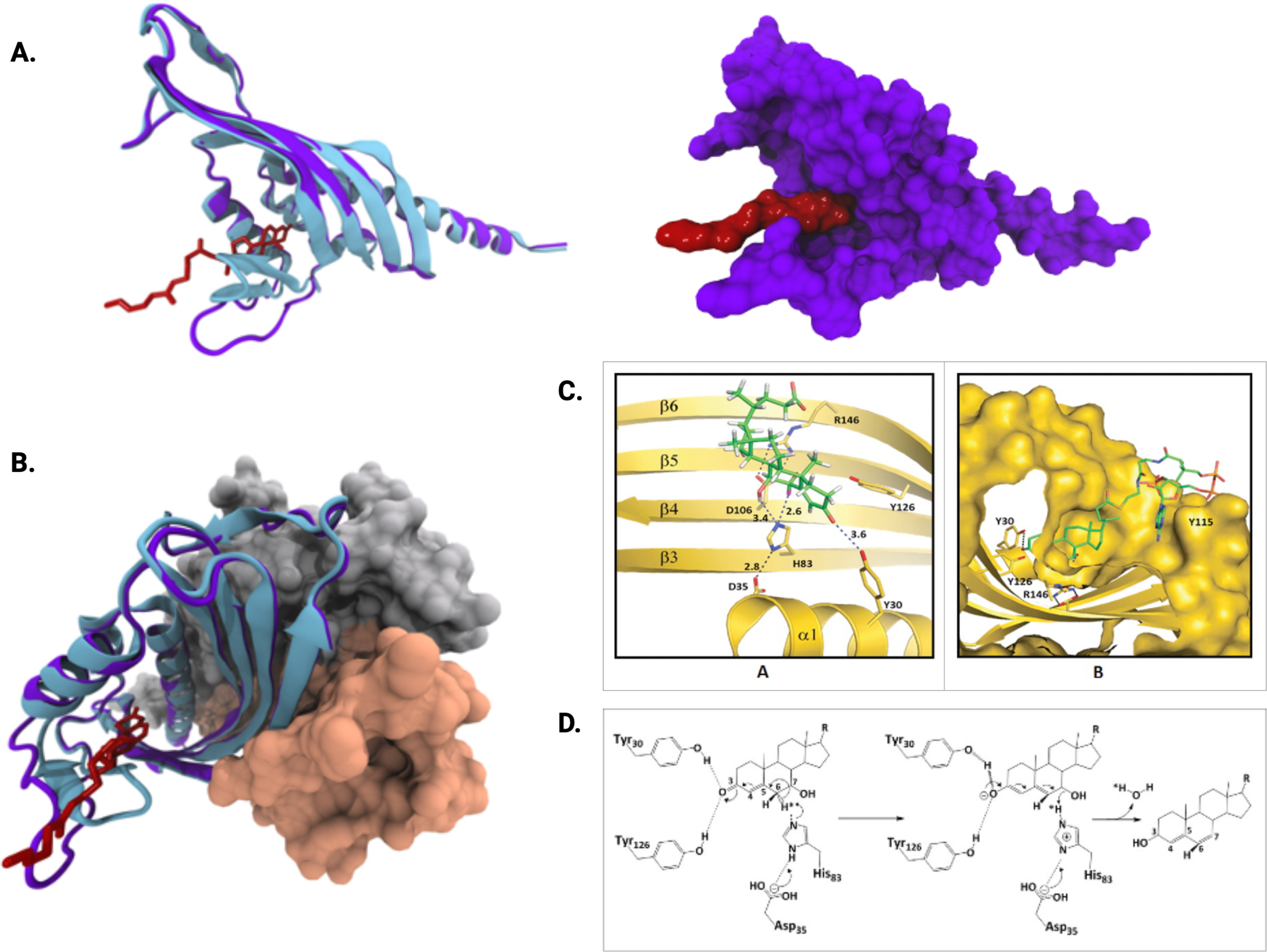
Gene organization of bile acid and steroid-metabolizing genes in *C.* *scindens* VPI12708 and *C. scindens* ATCC 35704. The complete genomes from each strain have been deposited previously ().

### BaiG: proton-dependent bile acid transporter

Bile acid 7ɑ-dehydroxylation in whole cells of *C. scindens* VPI 12708 occurs rapidly (White et al. 1980), yet bile acid intermediates do not appreciably accumulate intracellularly (White et al. 1981) indicating bile acid transport. Within the polycistronic *bai* operon is a 1.4-kb open reading frame encoding a 49.9-kDa polypeptide designated as *baiG*. The BaiG is annotated as a member of the multiple facilitator superfamily (MFS), and hydropathy analysis predicts 14 membrane-spanning domains (Mallonee et al. 1996). Transport was observed to increase with decreasing pH. Proton ionophores, but not potassium ionophores, were reported to inhibit bile acid transport by recombinant BaiG in *E. coli* indicating symport of bile acids driven by proton motive force (Mallonee et al. 1996). Additional kinetic and substrate specificity studies of *baiG* will be important in order to understand the relative rate of bile acid 7ɑ-dehydroxylation between bile acid substrates with whole cells of *C. scindens*. Recent studies in which *bai* genes were engineered into the chromosomes of *Clostridium sporogenes* (Funabashi et al. 2020) or *Escherichia coli* (Meibom et al. 2024) have also introduced the *baiG* to enhance transport, although future studies should examine whether *baiG* is required for efficient import of primary bile acids into the cell.

### BaiB, BaiF, and BaiK: Bile acid coenzyme A metabolism

Within the polycistronic *bai* operon in *C. scindens* strains are two genes encoding enzymes predicted to function in coenzyme A metabolism. The *baiB* gene is the first structural gene in the *bai* operon and was demonstrated to encode a 58-kDa ATP-dependent bile acid CoA ligase that catalyzes the first enzymatic step in the pathway leading to DCA (Mallonee et al. 1992). A crystal structure at 2.19 Å has been deposited for BaiB (PDB 4LGC) and awaits additional biochemical characterization. The *baiF* gene encodes a 47-kDa polypeptide predicted to encode a CoA hydrolase (Ye et al. 1999). The purified recombinant BaiF was predicted to be a dimer (72 kDa) by gel filtration with an apparent *Km* value 175 μM against cholyl∼CoA, indicating that primary bile acid CoA conjugates are likely not the physiological substrate (Ye et al. 1999). BaiF did not hydrolyze acetyl∼CoA, isovaleryl∼CoA, palmitoyl∼CoA, or phenylacetyl∼CoA (Ye et al. 1999), although it hasn’t been determined if substrates other than bile acids can accept CoA from bile acid∼CoA intermediates. Subsequently, recombinant BaiF was demonstrated to transfer CoA from deoxycholyl∼CoA, lithocholyl∼CoA, and allodeoxycholyl∼CoA to primary bile acids where CA > alloCA > β-murocholic acid (β-MCA) > ursodeoxycholic acid (UDCA) > chenodeoxycholic acid (CDCA) (Ridlon et al. 2012). Funabashi et al. (2020) reported that BaiF was not required for the rate limiting 7ɑ-dehydration step of CA in a stepwise pathway Bai enzyme reconstruction assay, as 3-oxo-4,6-DCA∼SCoA as well as 3-oxo-4,6-DCA accumulated in the presence of BaiB, BaiA2, BaiCD, and BaiE (BaiF and BaiH were left out). A recent study on CDCA conversion to LCA reported that only BaiB was needed for LCA formation, although CoA intermediates were found to accumulate (Meibom et al. 2024). BaiF is a member of the Type III CoA transferase family with conserved active-site D169 predicted to be involved in aspartyl∼CoA thioester formation, thereby releasing the bile acid pathway intermediate followed by regeneration of D169 via transfer to BaiG-transported primary bile acid (Ridlon et al. 2012).

We previously characterized a polycistronic *bai* operon in *Clostridium hylemonae* through genome-walking by PCR and determined that *baiA2* was not present in this operon (Ridlon et al. 2010). Using *baiA* nucleotide sequences from *baiA* genes from *C. scindens*, we amplified the partial *baiA1* from *C. hylemonae* using degenerate primers (Ridlon et al. 2010). Genome-walking in both directions from *baiA*1 resulted in identification of a novel gene cluster encoding a *baiF* homolog that we named *baiK* (Ridlon et al. 2012). The gene cluster was also located flanking the *baiA*1 gene in *C. scindens* VPI 12708, but not *C. scindens* ATCC 35704 (Ridlon et al. 2012). The *baiJ* genes encode a predicted 62-kDa flavoprotein similar to 3-ketosteroid-delta1-dehydrogenases. The *baiK* genes encode a predicted 49-kDa type III CoA transferase homologous to the *baiF* gene (63% amino acid identity) (Ridlon et al. 2012). The *baiL* genes are predicted to encode a 27-kDa protein in the SDR family. In *C. scindens* VPI 12708, we located a TspO/MBR family protein encoding gene downstream from *baiA1* but on the opposite strand (Ridlon et al. 2012). In *C. scindens* ATCC 35704, the TspO/MBR gene is downstream on the same strand and is bile acid-inducible (Devendran et al. 2019). We recently reported that phage-induced disruption of TspO expression in *Bacteroides vulgatus* reduced bile salt deconjugation (Campbell et al. 2020). Evidence that *baiJKL* operon is involved in bile salt metabolism was provided by demonstrating that recombinant BaiK catalyzed bile acid CoA transferase activity (Ridlon et al. 2012). The *baiJ* and *baiL* genes were also shown to be bile acid inducible and transcriptionally linked to *baiK* expression (Ridlon et al. 2012).

The rate-limiting bile acid 7-dehydration catalyzed by BaiE appears to recognize both free bile acid substrates as well as SCoA conjugates (Dawson et al. 1996). This is an energy conserving reaction and therefore important to understand. Proper kinetic analysis of BaiF and BaiK as well as genetic knock out of these genes detect is needed to determine the importance of these gene products in CoA metabolism as well as substrate specificity in vivo.

### BaiA: bile acid 3α-hydroxysteroid dehydrogenases

Several 27-kDa polypeptides appeared on denaturing 2D-gel following induction of cultures of *C. scindens* VPI 12708 (White et al. 1981). The *baiA2* was located within the large *bai* polycistronic operon (Mallonee et al. 1990), sharing 92% amino acid sequence identity with deduced amino acid sequences from monocistronic copies *baiA1* and *baiA3*, which are identical at the nucleotide level (Gopal-Srivastava et al. 1990). The *baiA1* was subsequently cloned and overexpressed in *E. coli* (Mallonee et al. 1995). The amino acid sequence indicated that BaiA proteins are members of the short chain reductase/dehydrogenase family of proteins that include hydroxysteroid dehydrogenases. The partially purified recombinant BaiA1, purified native BaiB, [24-^14^C] CA, ATP, NAD^+^ or NADP^+^, and coenzyme A yielded a product consistent with [24-^14^C] 3-oxo-cholyl∼CoA (Mallonee et al. 1995). Kinetic analysis of purified recombinant BaiA1 with either cholyl∼CoA or deoxycholyl∼CoA and pyridine nucleotide revealed that coenzyme A conjugates are preferred substrates as activity was not detected with unconjugated CA and DCA (Mallonee et al. 1995). These results are consistent with coenzyme A metabolism catalyzed by BaiB (ATP-dependent) and BaiF/BaiK (ATP-independent) described above.

Both the apo (1.9 Å) and NAD(H) bound (2.0 Å) crystal structures of tetrameric BaiA2 from *C. scindens* VPI 12708 were reported along with steady state kinetic analysis with both unconjugated primary and secondary bile acids, glycine and taurine conjugates, as well as coenzyme A conjugates of primary and secondary bile acids (Bhowmik et al. 2014). Steady state kinetics indicated that NAD+ is the preferred cofactor, and the binary structure revealed steric and electrostatic hindrance of the 2’-phosphate on NADP+. Indeed, the E42A mutant showed improved utilization of NADP+ (Bhowmik et al. 2014). Catalytic efficiency between unconjugated primary and secondary bile acids was two orders of magnitude lower than for the coenzyme A conjugates (Bhowmik et al. 2014). Recognition of both cholyl∼CoA and deoxycholyl∼CoA also indicates that BaiA1 and BaiA2 may act in both the first and final redox steps in the pathway (Bhowmik et al. 2014). Interestingly, transcriptomic analysis of *C. scindens* ATCC 35704 induced with CA resulted in significant up-regulation of *baiA1* and *baiA2*; however, induction with DCA resulted in downregulation of the *baiBCDEA2FGHI* operon, but upregulation of *baiA1* (Devendran et al. 2019). Recent work combining in vitro heterologous expression of *baiB, baiCD, baiE, baiA2, baiF, baiH,* and *baiI* with integration of *baiBCDEA2FHI* in *C. sporogenes*, which lacks the pathway, demonstrated that these genes were both necessary and sufficient to convert CA to DCA (Funabashi et al. 2020). BaiA2 (or BaiA1) was found to be sufficient for the first and last redox step in the formation of LCA from CDCA (Meibom et al. 2024). Future genetic studies will be needed to determine the relative roles of *baiA1* and *baiA2* in *C. scindens*.

### BaiCD and BaiH: oxidoreductases that differentiate between 7-hydroxy epimers

Early speculations about the source of reducing equivalents utilized in bile acid biotransformations by *C. scindens* were hinted at by NADH-dependent flavin oxidoreductase activity (NADH:FOR) that provides reduced flavins for the 21-dehydroxylation of deoxycorticosterone (Feighner et al. 1979). This prompted Lipsky and Hylemon to partially purify NADH:FOR from *C. scindens* VPI 12708 (Lipsky et al. 1980). Interestingly, the NADH:FOR that was characterized was shown to be induced by CA but not DCA (Lipsky et al. 1980). In 1993, Franklund and colleagues purified the native NADH:FOR 372-fold to apparent electrophoretic homogeneity with subunit and native molecular weight estimates of 72 kDa and 210 kDa, respectively (Franklund et al. 1993). The N-terminus of the polypeptide was sequenced, and an oligonucleotide was synthesized, allowing the gene to be mapped on the *bai* operon (Franklund et al. 1993). The gene was named “*baiH*”, and multiple-sequence alignment against characterized homologs indicated this polypeptide contains a conserved Fe-S center and flavin-binding site. Soon after, Baron and Hylemon cloned and heterologously expressed the *baiH* gene in *E. coli* (Baron et al. 1995). Each subunit of the purified recombinant BaiH contained 2 mol iron, 1 mol copper, and 1 mol FAD. It was determined during this work that the BaiH and BaiCD were paralogs and may catalyze a similar reaction, at the time, maintaining the cellular ratio of NAD/NADH.

As the oxidative branch of the pathway became clearer, there were two steps involving oxidation of the *C4-C5* in both CA/CDCA (7ɑ-hydroxy) and UDCA (7β-hydroxy) prior to the rate-limiting bile acid 7ɑ-dehydration (BaiE) and 7β-dehydration (BaiI?), respectively. The hypothesis was tested that *baiCD* and *baiH* encode stereospecific enzymes catalyzing oxidation of *C4-C5* of 3-oxo-CDCA or 3-oxo-UDCA by detecting product formation after TLC and LC/MS following incubation with each recombinant enzyme (Kang et al. 2008). It was determined that the *baiCD* gene encodes a stereo-specific NAD(H)-dependent 7ɑ-hydroxy-3-oxo-Δ^4^-cholenoic acid oxidoreductase, and the *baiH* gene encodes a stereospecific NAD(H)-dependent 7β-hydroxy-3-oxo-Δ^4^-cholenoic acid oxidoreductase (Kang et al. 2008).

Subsequent work determined that *baiH* and *baiCD* gene products also function in the reductive arm of the Hylemon-Björkhem Pathway during the conversion of CA to DCA (Funabashi et al. 2020). The BaiH functions as the elusive Δ^6^-reductase of Samuelsson and Bergström, but whose substrate is 3-dehydro-4,6-deoxycholate and/or 3-dehydro-4,6-deoxycholyl∼ScoA (Ridlon et al. 2023). There is clear economy in this pathway as *baiA*, *baiCD,* and in cases, *baiH* function in two separate steps with analogous substrates, thus reducing the number of genes required.

### The structure and catalytic mechanism of the rate-limiting bile acid 7ɑ-dehydratase encoded by the *baiE* gene

In 1981, the results of one- and two-dimensional SDS-PAGE of CA-induced vs. uninduced cell extracts from *C. scindens* VPI 12708 indicated the formation of at least five induced polypeptides, including one estimated at *M_r_* 23.5 kDa (White et al. 1981). Cloning and nucleotide sequencing of the *baiBCDEAF* genes followed by purification and N-terminal sequencing of the 23.5-kDa polypeptide resulted in identifying this polypeptide as the product of the *baiE* gene (deduced *M_r_* = 19.5 kDa), although the function was not known (Mallonee et al. 1990). Around this time, intermediates in the complex biochemical pathway resulting in conversion of CA to DCA were identified and determined by a collaborative effort between the microbiologist, Phillip Hylemon, and the bile acid chemist, Ingemar Bjӧrkhem (see **Figure 1** for their photos) (Hylemon et al. 1991). Thus, it was known that the substrate for the bile acid 7ɑ-dehydratase derived from CA is 7ɑ-,12ɑ-dihydroxyl-3-dehydro-4-cholenoic acid and the product is 12ɑ-hydroxy-3-dehydro-4,6-choldienoic acid (Hylemon et al. 1991).

In 1996, it was reported that the *baiE* gene product was purified after heterologous expression in *E. coli* and demonstrated to encode the bile acid 7ɑ-dehydratase (Dawson et al. 1996). The BaiE shares few primary sequence homologs, and early attempts to identify homologs began with collaborative efforts between Phillip Hylemon and Alexey Murzin of the MRC Laboratory of Molecular Biology at Cambridge University in the early 2000s (Ridlon et al. 2006). The computational model was based on secondary structural alignments between BaiE and scylatone dehydratase, nuclear transport factor 2, and steroid Δ^5^-isomerase, whose structures had been solved. The substrate, 7ɑ-,12ɑ-dihydroxyl-3-dehydro-4-cholenoic acid, was modeled into the active-site and the model originally reported in 2006 was confirmed following crystallization by Scott A. Lesley’s laboratory at the Scripps Institute, and site-directed mutagenesis of predicted active-site amino acids by the Hylemon lab (Bhowmik et al. 2016).

The BaiE structure from *C. scindens*, *C. hylemonae*, and *Clostridium hiranonis* (renamed *Peptacetobacter hiranonis* in 2020) were coupled with size-exclusion chromatography revealing trimeric quaternary structure, the monomers are composed of a single domain with characteristic ɑ + β barrel fold of the nuclear transcription factor 2-like superfamily of proteins and are linked together partly via divalent ion-His coordination (**Figure 6A-B**) (Bhowmik et al. 2016). A co-crystal between BaiE and the product 3-dehydro-4,6-lithocholyl∼CoA confirmed simulated docking experiments indicating substrate interactions with catalytic residues Y30, H83, R146, Y126, and D106 (**Figure 6C**). The coenzyme A moiety is presumed to extend into bulk solvent but appears to be important in catalysis with ∼10-fold higher catalytic efficiency. The co-crystal also revealed a novel extended pocket which was not predicted in the computational model in which a loop (residues 48-63) forms the extended pocket. Based on active-site architecture and site-directed mutagenesis data, a catalytic mechanism has been proposed. Y30 acts as a general acid (assisted by Y126) facilitating the delocalization of π-electrons between *C3, C4, C5,* and *C6*. Y30 is predicted to protonate the oxyanion on the bile acid *C3*-oxo group, stabilizing the negative charge. H83 is positioned to abstract the destabilized 6ɑH and protonate the leaving *C7*-hydroxyl group. D35 is important for maintaining the P*K*_a_ of H83 ensuring the release of a water molecule (**Figure 6D**) (Bhowmik et al. 2016).

**Figure 6.**
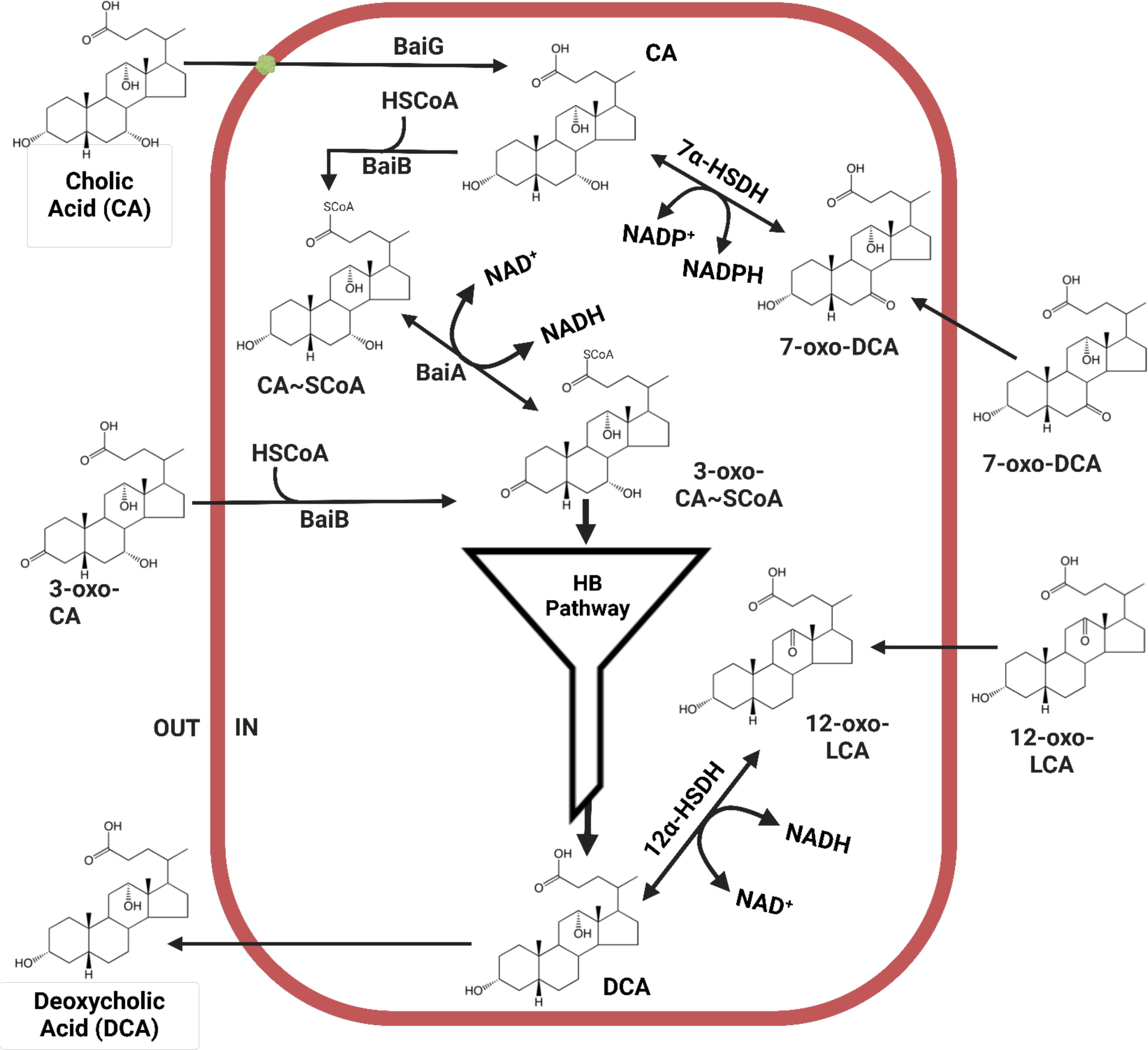
Structure and catalytic mechanism of BaiE, the bile acid 7a-dehydratase. **A.** Visual Molecular Dynamics (VMD) Model of BaiE, a potential drug target, and rate-limiting enzyme responsible for the formation of toxic and cancer-causing bile acids. Left: Ribbon diagram of Ligand-bound subunit (purple) is overlaid with apo-enzyme (cyan). Bile acid ligand displayed in red. Right: Monomeric space-filling subunit structure of BaiE from *C. hiranonis* (purple) with bile acid ligand (red). **B.** Trimeric native form of BaiE from *C. hiranonis* with mixed ribbon and space-filling subunits. Ligand-bound subunit (purple) is overlaid with apo-enzyme (cyan). Bile acid ligand displayed in red. **C.** Left: Probable productive binding mode of 3-oxo-Δ^4^-CDCA. Blue dashed lines and adjacent numbers are predicted interaction of His83 with C7-OH and C6 atoms and Y30-OH group with *C3*-oxo atom of 3-oxo-Δ^4^-CDCA. The 6α-H closest to H83 colored magenta and 6β-H away from H83-Nε2 atom colored brown. Right: Predicted stacking interaction involving the adenine group of the Coenzyme (CoA) moiety of 3-oxo-Δ^4^-CDCA∼SCoA with Y115. The key interaction of the bile acid moiety of the docked CoA-bile acid ester with the active site residues is like what is predicted in left panel. Carbon atoms of protein residues and product molecule are colored gold and green, respectively. H, O, N, P and S atoms are colored gray, red, blue, orange and olive, respectively. **D.** Proposed mechanism of catalysis by BaiE. Y30 acts as a general acid protonating *C3*-oxyanion, stabilizing negative charge, and potentiating electron shift, destabilizing *C6*-6αH. H83, stabilized by D35, acts as a general base, executing deprotonation and ensuring protonation reaction with the subsequent release of water. Figure modified from previously published images with kind permission ().

The *baiE* gene represents a key target for gene knockout of 7ɑ-dehydroxylating activity against CDCA and CA since this represents the only irreversible step in the pathway following transport. So far, a genetic system for *C. scindens* has not been reported. The BaiE may also represent a drug target to inhibit the formation of hydrophobic bile acids DCA and LCA, particularly since bacteria encoding the *bai* pathway are rare in the gut microbiome.

### Bile acid 7β-dehydratase

Whole cells and cell-free extracts of *C. scindens* have been shown previously to catalyze the conversion of the 7β-hydroxy bile acid ursodeoxycholic acid (UDCA) to LCA (White et al. 1982). The addition of NAD+ stimulated activity in cell extracts, and 7β-dehydratase activity against UDCA was CA-inducible (White et al. 1982). In this study, both 7ɑ- and 7β-dehydratase activities were inactivated by heating to 45°C, and both co-eluted in a single peak at 114 kDa. This suggested that either the enzymes are of the same size and stability, or that a single enzyme recognizes both stereo-chemistries. However, when the *baiE* was later expressed and both stereo-chemistries were tested, it was clear that BaiE did not recognize 7β-hydroxylated substrates (Dawson et al. 1996). In addition, the oxidation step prior to 7ɑ- and 7β-dehydroxylation required two stereospecific enzymes encoded by *baiCD* (7ɑ-hydroxy specific) and *baiH* (7β-hydroxy specific) genes (Kang et al. 2008). Indeed, at the time the BaiE was first characterized, the *baiBCDEAFGHI* operon had been cloned and sequenced and the deduced amino acid sequence of *baiI* indicate that this protein shares both the same SnoaL_4 protein superfamily and subunit *M_r_* which if this enzyme exists as a trimer, would explain the co-elution observed in an earlier study (White et al. 1982). Bile acid 7ɑ- and 7β-dehydroxylating activities are both induced by CA and to a lesser extent CDCA, but not UDCA. This is also true of bile acid induction of the *baiBCDEAFGHI* polycistronic mRNA (White et al. 1988).

An alternative hypothesis that might be advanced is that a 7β-dehydratase is not necessary for the 7β-dehydroxylation of UDCA since *C. scindens* encodes NADP-dependent 7ɑ-HSDH, and provided that this bacterium also encode NAD(P)-dependent 7β-HSDH. In this scheme, UDCA could be oxidized to 7-dehydro-LCA and reduced to CDCA. CDCA could then be 7ɑ-dehydroxylated by the *bai* operon, including the rate-limiting 7ɑ-dehydration by BaiE. Numerous studies that have examined in vitro bile acid metabolism of CDCA and CA by whole cells and cell-free extracts and have not identified detectable accumulation of 7β-hydroxylated intermediates. This suggests that *C. scindens* does not encode NAD(P)-dependent 7β-HSDH. Taken together, the more parsimonious explanation these data suggest is that the *baiI* gene encodes a bile acid 7β-dehydratase, although this has not been demonstrated empirically to date.

### Flavoproteins involved in the “reductive arm” of DCA production

The removal of the 7ɑ/β-hydroxyl group results in formation of a stable 3-dehydro-4,6-choldienoic acid intermediate (Hylemon et al. 1991). Three sequential reductions have been hypothesized, requiring flavoproteins for reduction of ring A (C_4_-C_5_) and ring B (C_6_-C_7_) in addition to pyridine nucleotide-dependent 3ɑ-HSDH. In support of this, reduced flavins stimulated bile acid 7α-dehydroxylation in cell-free extracts of *C. scindens* VPI 12708 (White et al. 1983). While work on the *bai* regulon progressed from the 1980s to 2000s allowing more detailed understanding of oxidative steps in the pathway from bile acid transport (*baiG*) to the rate-limiting 7ɑ-dehydration step (*baiE*), progress on the reductive arm of the pathway has only been made recently. In 2018, a flavoprotein was identified among a list of flavin-dependent enzymes in the genome of *C. scindens* ATCC 35704 that was annotated as a flavin-dependent “squalene desaturase”, involved in binding a precursor of cholesterol biosynthesis (Harris et al. 2018). A homolog of this gene was also identified in all strains of *C. scindens* characterized to date (Caicedo et al. 2024) and other bile acid 7ɑ-dehydroxylating bacteria indicating that this is a candidate 5β-reductase.

Incubation of the purified recombinant 45.4-kDa flavoprotein with 3-dehydro-DCA under aerobic conditions resulted in formation of product irrespective of pyridine nucleotide addition (Harris et al. 2018). This is suspected to be due to auto-oxidation of the bound flavin. However, the protein was relatively unstable and precipitates after only a few hours following affinity purification. Another study concluded that BaiN was not required for conversion of CDCA to LCA; however, it is not clear that the purified enzyme was active when applied to the multi-enzyme in vitro assay (Meibom et al. 2024). In our study, the enzyme-catalyzed reaction product was observed from multiple enzyme preparations both in the Ridlon lab at the University of Illinois and the Hylemon lab at Virginia Commonwealth University. The product was subjected to LC/MS-IT-TOF analysis, and we expected a loss of two amu but observed a loss of four. This appears to indicate that a single enzyme may be sufficient for conversion of 3-dehydro-4,6-DCA (product of BaiE) to 3-dehydro-4-DCA and then to 3-dehydro-DCA (Harris et al. 2018). These same reactions were shown to be catalyzed by BaiH and BaiCD (Funabashi et al. 2020). A study by Meibom et al. (2024) indicates that the BaiP (they refer to as BaiO) (but not the BaiJ) also catalyzes a two-step reduction from 3-dehydro-4,6-LCA to 3-dehydro-4-LCA and then to 3-dehydro-LCA (**Figure 4**). The relative contribution of each of these Bai reductases in the metabolism of primary to secondary bile acids requires additional work.

The BaiN sequence appears to be conserved in *Clostridium* spp., and the *baiN* gene product has been demonstrated to convert CA to DCA. Furthermore, sequence ID decreases significantly in non-DCA forming *Firmicutes* indicating that this enzyme is likely specialized for bile acid 7ɑ-dehydroxylation. The Hylemon-Bjorkhem model for bile acid 7ɑ-dehydroxylation was based on the accumulation of radiolabeled CA intermediates extracted after incubation with cell extracts of *C. scindens* VPI 12708 (Hylemon et al. 1991). In this study, 3-dehydro-4-DCA and 3-dehydro4,6-DCA intermediates were detected, which co-migrated, and must be separated by argentation chromatography. It was hypothesized that two flavoproteins would be necessary to catalyze *C4-C5* followed by *C6-C7* reduction (Hylemon et al. 1991). Further studies will be needed to determine the kinetics and substrate-specificity of the *baiN* gene product.

A recent approach of combining recombinant *baiBCDEAFH* enzymes in vitro and engineering the *bai* operon into the chromosome of *C. sporogenes* suggests that the *baiBCDEAFGH* genes are needed for conversion of CA to DCA (Funabashi et al. 2020). Taken together, in the case of conversion of CA to DCA, the *baiCD* functions in both the second oxidative and second to last reductive step, and the *baiH* function in the first reductive step in the pathway. Further research is needed in order to determine the relative contribution of flavoproteins encoded by *baiN*, *baiCD*, and *baiH* to the reductive and oxidative arms of the pathway.

### Final enzymatic steps and secondary bile acid export

Following reduction of C_4_-C_5_ and C_6_-C_7_ the 3-keto group is reduced, and the bile acid exported from the cell. There is still uncertainty regarding the point in the pathway in which the BaiF and BaiK transfer CoA. There is reason to think that CoA-transfer occurs after the rate-limiting 7ɑ-dehydration catalyzed by BaiE or BaiI. The BaiE recognizes substrates irrespective of CoA conjugation, the CoA moiety protrudes from the active site into bulk solvent (Bhowmik et al. 2014). Earlier studies indicated that 3-oxo-4-DCA, a metabolite downstream from 7-dehydration, is linked to what appears to be CoA (Coleman et al. 1987). Identifying the major metabolite(s) that are CoA-conjugated may be settled by LC/MS analysis at various time points of quenched cell extracts from CA-induced *C. scindens* whole cells following addition of CA.

The final oxidation step, conversion of 3-oxo-DCA(∼SCoA) or 3-oxo-LCA(∼SCoA) to DCA(∼SCoA) or LCA(∼SCoA), is expected to proceed via an NAD(P)H-dependent HSDH. A recent study (Heinken et al. 2019) of bile acid-metabolizing genes in stool metagenomes from patients with inflammatory bowel disease versus healthy age-matched control stool samples suggested a candidate gene for this step in the pathway, naming it the *baiO* (CLOSCI_00522). The gene itself encodes a putative 62-kDa flavoprotein which is directly downstream of the *baiN* that we recently reported (Harris et al. 2018) encodes a flavoprotein involved in reduction of C_4_-C_5_ and C_6_-C_7_. Biochemical characterization of the 62-kDa flavoprotein has not been reported, but the rationale for naming this gene *baiO* is that bacteria often cluster genes involved in particular pathways together, and because this gene is clustered with the *baiN* and based on annotation utilizes pyridine nucleotide, it is probable that this enzyme catalyzes the only oxidative step for which there is yet no known enzyme (Heinken et al. 2019). So far, bacterial HSDHs are found in the short or medium chain dehydrogenase families as well as the aldo-keto reductase family which do not utilize flavins. These enzymes range in subunit size typically from 25 to 37 kDa. While this does not exclude the predicted *baiO*, it would suggest this gene would be an exception to the rule in terms of genes that gut microbes have evolved to metabolize diverse bile acid and steroid hydroxyl/carbonyl groups.

The most probable candidate for the final reductive step is that one or both copies of the *baiA* (*baiA1* and *baiA2*) act in both the oxidative arm and the reductive arm. Kinetic analysis of BaiA1 and BaiA2 suggest this is the likely scenario as the enzymes recognized 3-oxo-CA∼CoA and 3-oxo-DCA∼CoA to about the same extent. In our recent in vitro transcriptome analysis of *C. scindens* ATCC 35704 (Devendran et al. 2019), we reported that CA led to the induction of both copies of *baiA*; however, DCA addition significantly down-regulated the *baiBCDEAFGHI* operon, but the *baiA1* gene was significantly up-regulated by DCA. This may indicate that the *baiA1* gene is important in the last reductive step of the pathway.

The export of toxic secondary bile acids is likely facilitated by an ABC transporter such as a multi-drug efflux pump. A separate argument was made for the identity of the bile acid efflux pump (Devendran et al. 2019, Heinken et al. 2019). In this study, the reasoning behind searching for a shared export protein between *C. scindens* and *Eggerthella lenta* is based on the assumption that both encode *bai* operons and produce DCA. Putative *baiN* and *baiO* orthologs (*Elen_1017* and *1018*) were identified by BLAST, sharing 32% and 45% amino acid identity, respectively. Upstream of these genes is a putative transport protein (*Elen_1016*), which when BLAST searched against the *C. scindens* genome revealed an ortholog (*CLOSCI_01264*) that shared 59% protein identity. This gene was provisionally named the *baiP* on the basis of this sequence comparison alone (Heinken et al. 2019). One difficulty with this hypothesis is that *E. lenta* encodes a “*bai-like operon*” and has been repeatedly shown to oxidize and epimerize bile acid hydroxyl groups, but lacks bile acid 7ɑ-dehydroxylating activity. Currently, the export of secondary bile acids has yet to be characterized.

## Formation of allo-secondary bile acids by *C. scindens*

While the adult human liver generates two primary bile acids, CDCA and CA, there is great diversity in bile acid structure among vertebrates (Hofmann et al. 2010). Nine [24-^14^C] CA intermediates were identified after incubation with cell-free extracts of CA-induced whole cells of *C. scindens* VPI 12708 (Hylemon et al. 1991). Each metabolite was identified by characterization with stereospecific 3ɑ-, 7ɑ-, 12ɑ-, and 3β-HSDH enzymes, relative migration on TLC and HPLC against known bile acid standards, and GLC/MS analysis (Hylemon et al. 1991). Two unknown metabolites were identified. Each metabolite had similar, but not identical migration with DCA and 3-dehydroDCA, respectively. Since the HSDH panel showed identical patterns with DCA and 3-dehydroDCA, GLC-MS was required to identify the compounds in question. GLC retention time and mass spectra of the unknown compounds was identical to allo-DCA and allo-3-dehydro-DCA, respectively. Once identified, the [24-^14^C] CA metabolites were then chemically synthesized, individually added to cell extracts of CA-induced whole cells of *C. scindens* VPI 12708 and shown to be converted to DCA or allo-DCA (Hylemon et al. 1991). Thus, while hepatocytes are capable of generating primary allo-bile acids (e.g., allo-CA and allo-CDCA) (Shiffka et al. 2017, Shiffka et al. 2020), allo-secondary bile acids appear to be end-products of microbial bile acid 7ɑ-dehydroxylation. Whether hepatocytes can convert DCA to alloDCA has not been addressed to our knowledge.

[24-^14^C] alloDCA is formed in cell-free extracts of *C. scindens* VPI 12708 on the order of 4 micromolar (Hylemon et al. 1991), but typically when whole cells of *C. scindens* strains are induced and bile acids extracted from the spent medium, conversion to alloDCA is minimal if observed at all. A defined medium recently developed for the cultivation of *C. scindens* ATCC 35704 has been used to assess the transcriptional profiles to the bile acids CA and DCA (Devendran et al. 2019). One of the observations was induction 3.82 log_2_FC (FDR = 5.35E-26) by CA, but not DCA, of an uncharacterized flavoprotein (HDCHBGLK_03451). In a subsequent study (Lee et al. 2022), HDCHBGLK_03451 was cloned and the recombinant enzyme expressed in *E. coli*. It was determined that resting cells expressing HDCHBGLK_03451 yielded the bile acid product 3-dehydro-alloDCA from 3-dehydro-4-DCA and 3-dehydro-alloLCA from 3-dehydro-4-LCA. When co-expressed with *baiA2*, 3-dehydro-4-DCA was converted to alloDCA and 3-dehydro-4-LCA was converted to alloLCA. We suggested the name *baiP* for HDCHBGLK_03451. Phylogenetic analysis of BaiP revealed a separate, but closely related gene cluster that contained the *baiJ* gene product, whose function was unknown, from *C. scindens* VPI 12708 and *C. hylemonae* TN271. We expressed the *baiJ* in *E. coli* and determined that BaiJ had bile acid 5ɑ-reductase activity similar to BaiP (Lee et al. 2022) (**Figures 4 and 5**). Thus, the remaining enzymatic steps in the Hylemon-Björkhem pathway involved in secondary allo-bile acid formation have been identified.

## Function of bile acid 7ɑ-HSDH and 12ɑ-HSDH

The reversible oxidation and reduction of bile acid hydroxyl groups is a phenotype harbored by diverse gut microbiota (Doden et al. 2021). Consequently, *C. scindens* must be capable of reducing oxidized bile acid hydroxyl groups (**Figure 7**). A constitutively expressed, native NADP(H)-dependent 7ɑ-HSDH was purified to electrophoretic homogeneity from *C. scindens* V.P.I. 12708 (Baron et al. 1991). The enzyme is a tetramer with a subunit mass of 32 kDa. The N-terminal sequence suggested that the enzyme was in the short chain dehydrogenase/reductase (SDR) family of enzymes. A reverse genetic approach was then used to synthesize a probe to locate and clone the gene using Southern blot. The gene encoding NADP-dependent 7ɑ-HSDH was cloned and overexpressed in *E. coli* and was shown to have similar subunit molecular mass, kinetic properties, and substrate specificity with the native enzyme (Baron et al. 1991). The -10 and -30 elements are distinct from conserved *bai* promoter region, and homologous to the constitutive promoter controlling expression of NAD-dependent 7ɑ-HSDH from *E. coli* (Baron et al. 1991). Indeed, 7ɑ-HSDH is widely encoded in diverse taxa in the gut environment, and 7-oxo-bile acid derivatives are detected in stool (Ridlon et al. 2006). The oxidized bile acid product (7-oxo-DCA) of CA is not a substrate for the BaiE (Dawson et al. 1996). The *bai* pathway oxidative steps utilize the NAD/NADH pool, while NADP-dependent 7ɑ-HSDH utilizes NADP/NADPH as co-substrate (White et al. 1983, Baron et al. 1991). Taken together, the NADP-dependent 7ɑ-HSDH appears to represent a regulatory pathway that can “switch” CA and CDCA “off” through oxidation and “on” through reduction, allowing the 7ɑ-hydroxyl group to be removed and the redox potential of the cell to be rapidly balanced.

**Figure 7.**
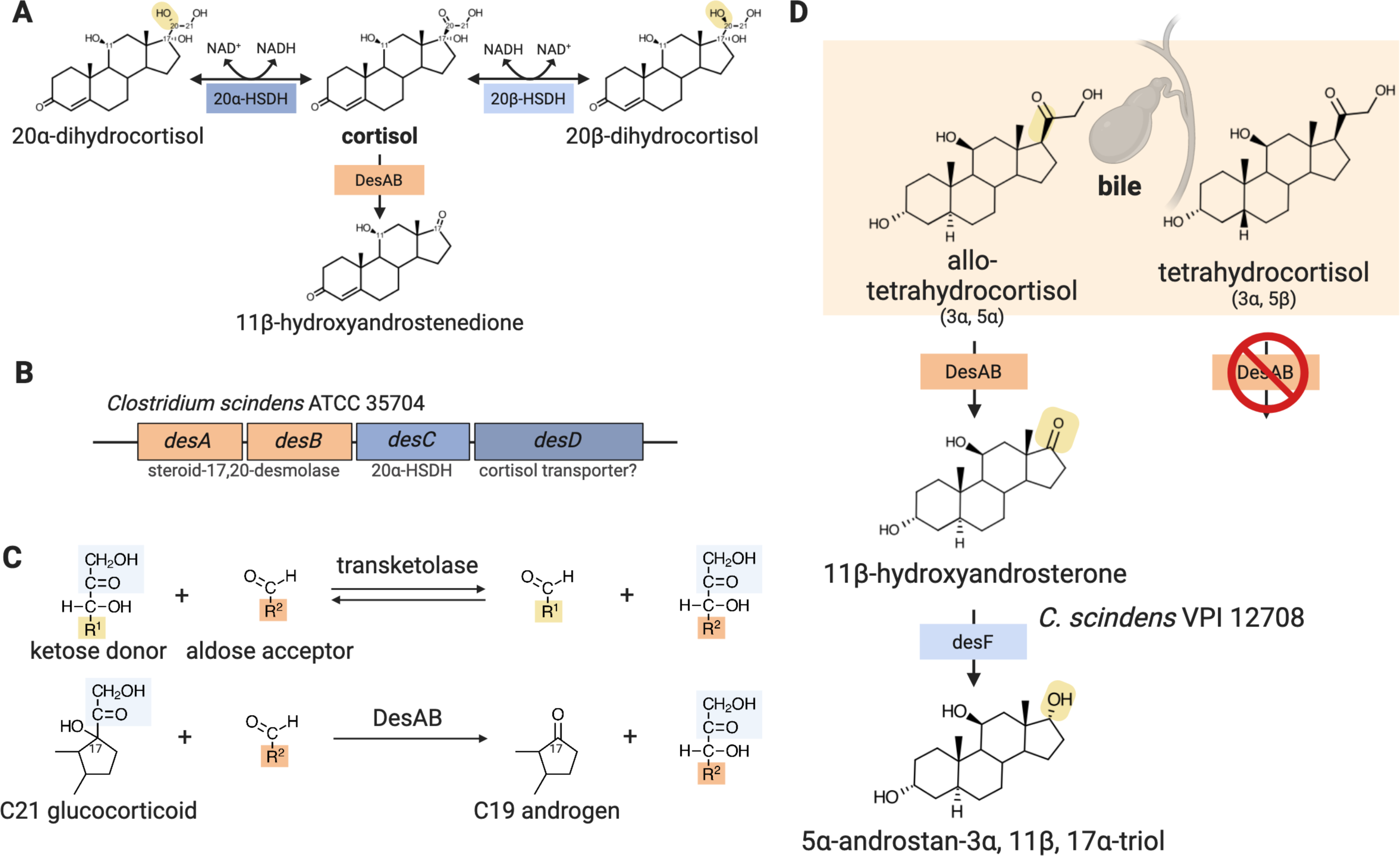
Bile acid oxidoreduction by *Clostridium scindens*. While CA is known to be transported by BaiG, it is assumed that oxo-bile acids are also recognized by this transporter, but this has yet to be determined. The import of 3-oxo-CA (or 3-oxo-CDCA) is predicted to be ligated to coenzyme A by BaiB and funneled into the HBP and converted to DCA. 7-oxo-DCA has been shown to be converted to CA by NADPH-dependent 7a-HSDH (). CA is then ligated to coenzyme A by BaiB and oxidized to 3-oxo-CA∼SCoA by the BaiA (NADH-dependent 3a-HSDH) (). 12-oxo-LCA is converted to DCA by NADH-dependent 12a-HSDH ().

An early study provided a major clue to the potential of bile acid 7ɑ-dehydroxylating bacteria, including *C. scindens*, to oxidize/dehydrogenate the 12ɑ-hydroxyl group of CA derivatives (Masuda et al. 1983). Aerobic incubation of whole cells of strain HD-17 (later named *C. hiranonis*) resulted in abolition of bile acid 7ɑ-dehydroxylation and instead conversion first to 7-oxo-DCA followed by formation of 7,12-dioxo-cholanoic acid (Masuda et al. 1983). A few years prior to this study, a 12ɑ-HSDH was partially purified and characterized from *Clostridium* group P C48-50 ATCC 29733 (Mahony et al. 1977, Macdonald et al. 1979). The nucleotide and amino acid sequence were reported in 2011 in a patent (Aigner et al. 2011). From this sequence, a gene was identified in *C. scindens*, *C. hylemonae*, and *C. hiranonis* based on phylogenetic analysis to the 12ɑ-HSDH from *Clostridium* group P C48-50 ATCC 29733 (Kisiela et al. 2012). Biochemical confirmation of 12ɑ-HSDH activity in bile acid 7ɑ-dehydroxylating bacteria was subsequently reported by our group (Doden et al. 2018). Recombinant 12ɑ-HSDHs displayed an order of magnitude lower activity towards 12-oxo-CDCA relative to 12-oxo-LCA (Doden et al. 2018). Catalytic efficiencies (*Km/Kcat*) were ∼3 fold greater in the reductive direction, with substrate-specificities. Marion et al. (2019) confirmed prior studies characterizing 12ɑ-HSDH activity in bile acid 7ɑ-dehydroxylating bacteria. Lysozyme-treated pellets of *C. scindens* ATCC 35704 incubated with CA resulted in accumulation of 12-oxo-lithocholic acid. Anaerobic cell-free extracts and whole cells rapidly reduced 12-oxo-LCA to DCA in vitro (Marion et al. 2019).

In addition, expressed recombinant proteins identified in the phylogeny were shown to have NADP-dependent 12ɑ-HSDH activity (Doden et al. 2018). Interestingly, substrate-specificity favored DCA over CA and the reductive direction. Our phylogenetic analysis and functional characterization of 12ɑ-HSDHs from *E. lenta* indicate that 12ɑ-HSDH activity is widespread in the gut microbiota and may favor the oxidative direction (Doden et al. 2018, Harris et al. 2018, Mythen et al. 2018). Ian MacDonald (see **Figure** 1 for his picture) was a pioneer in the study of HSDH enzymes from *E. lenta* (Macdonald et al. 1977, Macdonald 1978, Macdonald et al. 1979). Studies of bile acid hydrophobicity and toxicity indicate that 12-oxo-LCA is intermediate in hydrophobicity between CA and DCA as well as in toxicity towards Gram-negative bacteria (Watanabe et al. 2017). Collectively, this suggests that bile acid 7ɑ-dehydroxylating bacteria utilize 12ɑ-HSDH to maintain toxic concentrations of DCA. This is in contrast to other bacteria that have evolved the ability to oxidize DCA to 12-oxo-LCA in order to reduce toxicity. Genetic mutants of these enzymes and in vivo studies will be necessary to test this hypothesis in the future.

## Physiological responses of *C. scindens* to bile acids

While much of the work on *C. scindens* has been at the level of the *bai* regulon, recent work has sought to understand how bile acids affect global gene expression in *C. scindens* and host-microbe and microbe-microbe interactions in animal models. We recently performed RNA-Seq analysis of rRNA-depleted total RNA from *C. scindens* ATCC 35704 cultivated in our recently developed defined medium and compared this with defined medium supplemented with either 0.1 mM CA or 0.1 mM DCA (Devendran et al. 2019). We identified a total of 1,430 genes significantly differentially regulated by CA. There were 697 genes upregulated and 733 genes downregulated. DCA upregulated 684 genes and downregulated 1,033 genes. There were 897 genes shared between CA and DCA, while 278 were unique to CA and 207 unique to DCA. Clusters of orthologous groups altered by DCA included energy production and conservation (group C), and unknown function (group S), while CA altered group C and downregulated replication and repair (Group L) (Devendran et al. 2019). The *bai* genes were among the most highly expressed in the presence of CA, but downregulated significantly by DCA relative to control. One exception was the *baiA1* which was induced by DCA, perhaps suggesting this copy of the 3ɑ-HSDH is involved in the final oxidative step in the pathway leading to conversion of 3-dehydro-DCA to DCA (Bhowmik et al. 2014, Devendran et al. 2019).

We also recently characterized a novel isolate from pig feces designated *C. scindens* strain BL-389-WT-3D (DSM 100975) (Wylensek et al. 2020). The genome was sequenced and closed using a combination of Oxford nanopore and Illumina sequencing. Of the 3,655 predicted protein-encoding genes in DSM 100975, 2,966 genes were shared with *C. scindens* ATCC 35704. Genes that were unique to each strain appear to be composed largely of phage and mobile genetic elements, indicating the acquisition of distinct mobile elements unique to their respective host environments (Wylensek et al. 2020). Interestingly, the pig isolate did not grow in the defined medium developed for ATCC 35704, indicating additional growth requirements. We performed bile acid induction with CA and DCA in peptone yeast fructose (PYF) medium. The organization of the *bai* polycistronic operon in *C. scindens* DSM 100975 was nearly identical to human isolates ATCC 35704 and VPI 12708 except that there is a single ORF of unknown function inserted between *baiH* and *baiI* (Wylensek et al. 2020). RNA-Seq analysis indicates global transcriptional changes in the presence of bile acids (1,393 genes upregulated by CA, 1,336 downregulated), with significant upregulation of *bai* polycistronic genes by CA, but not DCA. The *baiN* gene in both ATCC 35704 and DSM 100975 was constitutive in the former but downregulated in the latter.

Bile acids induce expression of the well-studied multidrug efflux pump encoded by *acrAB* genes in *E. coli* (Rosenberg et al. 2003). It would be expected that the candidate bile acid efflux pump in *C. scindens,* which lacks *acrAB* homologs, would be induced by bile acids. Our recent transcriptome analysis of *C. scindens* ATCC 35704 identified several candidates including multidrug export permease *ygaD* homolog (HDCHBGLK_00878), ABC transporter *yxlF* (HDCHBGLK_01721), and multidrug resistance protein 3 (HDCHBGLK_02921) (Devendran et al. 2019). Future biochemical and/or genetic work will be required to determine the identity of the bile acid efflux pump(s) in *C. scindens*.

## Side-chain metabolism of cortisol

*C. scindens* ATCC 35704 was originally selected and isolated based on steroid-17,20-desmolase activity (Cerone-McLernon et al. 1981, Bokkenheuser et al. 1984, Morris et al. 1985). Whole cell and partially purified cell extract of *C. scindens* ATCC 35704 exhibited cortisol-inducible steroid-17,20-desmolase as well as NADH-dependent 20ɑ-HSDH activities (Krafft et al. 1987). Substrates for both steroid-17,20-desmolase and 20ɑ-HSDH were reported to have an absolute requirement for adrenocorticoids with 17ɑ,21-dihydroxy groups (Krafft et al. 1989).

In 2007, a draft genome of *C. scindens* ATCC 35704 became available on NCBI as part of the Human Microbiome Project (BioSample: SAMN00627066). Thereafter, we reported the first transcriptome analysis for *C. scindens* ATCC 35704 following cortisol-induction (Ridlon et al. 2013) (**Figure 2**). This approach resulted in identification of a gene cluster encoding steroid-17,20-desmolase (*desAB*) and NADH-dependent 20ɑ-HSDH (*desC*) as well as a putative cortisol transporter (*desD*) (**Figure 8**) (Ridlon et al. 2013, Devendran et al. 2018). Whereas aerobic mammals encode P450 monooxygenases involved in steroid side-chain cleavage (Bloem et al. 2013, Schiffer et al. 2019), anaerobic gut bacteria appear to utilize a novel oxygen-independent steroid transketolase enzyme encoded by *desAB* genes (Ridlon et al. 2013, Devendran et al. 2018).

**Figure 8.**
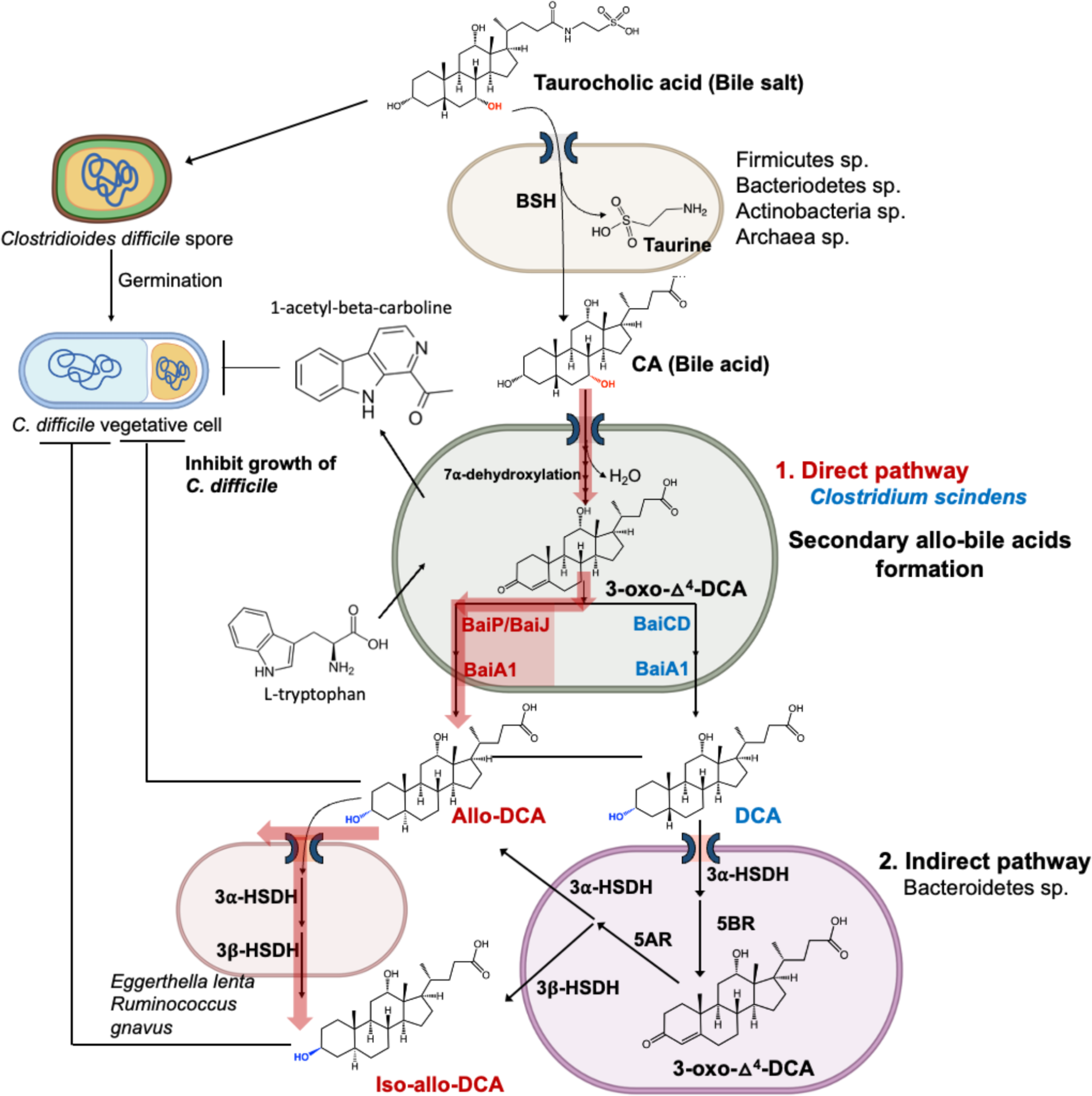
The steroid-17,20-desmolase pathway in host associated bacteria, including *C. scindens*. **A.** 20a-dihydrocortisol is converted to cortisol by DesC (NADH-dependent 20a-HSDH) and cortisol to 11β-hydroxyandrostenedione by DesAB (steroid-17,20-desmolase) encoded by *C. scindens* (). 20β-dihydrocortisol is not a substrate for *C. scindens*; however, organisms such as *Butyricicoccus desmolans, Clostridium cadavaris,* and *Propionimicrobium lymphophilum* express DesE, an NADH-dependent 20β-HSDH (). **B.** Gene cluster *desABCD* encoding steroid-17,20-desmolase (DesAB), DesC (NADH-dependent 20a-HSDH), and a putative cortisol transport protein (DesD) in *C. scindens* ATCC 35704 (). **C.** The *desA* and *desB* genes encode predicted N-terminal and C-terminal transketolases. An analogous reaction is predicted between sugar transketolation and steroid-17,20-desmolase. **D.** The host liver reduces cortisol to tetrahydrocortisol or allotetrahydrocortiol, some of which undergoes enterohepatic circulation via the bile. *C. scindens* is capable of recognizing allotetrahydrocortisol, converting this to 11β-hydroxyandrosterone. We recently discovered that *C. scindens* VPI 12708 encodes the *desF* gene which converts 17-keto androstanes to derivatives of epitestosterone. Epitestosterone and the 5a-reduced derivative of epiT (5a-dihydroepiT) has been shown to be an androgen receptor agonist (). Modified from previously published image with kind permission ().

Phylogenetic and sequence similarity networks based on the *desAB* genes in *C. scindens* ATCC 35704 resulted in identification of *desAB* genes in other taxa previously reported to express steroid-17,20-desmolase, namely *Butyricicoccus desmolans* (formerly *Eubacterium desmolans*) and *Clostridium cadaveris* (Bokkenheuser et al. 1986, Devendran et al. 2017, Ly et al. 2020). While *C. scindens* was reported to express cortisol 20ɑ-HSDH activity, which we demonstrated was encoded by *desC* gene (Ridlon et al. 2013), *B. desmolans* and *C. cadaveris* were reported to express cortisol 20β-HSDH activity (Bokkenheuser et al. 1986). We identified and characterized the gene (*desE*) encoding 20β-HSDH in *B. desmolans* and *C. cadaveris*, which is clustered with *desAB* (Devendran et al. 2017, Doden et al. 2019). Sequence-based analysis of *desAB* also revealed members of the urinary tract, such as *Propionibacterium lymphophilum*, encoding *desABE* (Ly et al. 2020). *P. lymphophilum* in the urinary has been shown to be correlated with prostate cancer (Shrestha et al. 2018). This unexpected observation led us to acquire, screen, and confirm metabolism of a range of endogenous and pharmaceutical glucocorticoids, several of which are therapeutic in prostate cancer such as prednisone and dexamethasone, by *P. lymphophilum* as well as *C. scindens* ATCC 35705 (Ly et al. 2020).

## Metabolism of androstanes

Screening of over a dozen strains of *C. scindens* for steroid-17,20-desmolase activity indicates that this function is rare in this species (Ridlon et al. 2013). Indeed, while *C. scindens* VPI 12708 lacks the ability to side-chain cleave cortisol, this strain expresses 17ɑ-HSDH which is predicted to convert the product of steroid-17,20-desmolase, 11β-hydroxyandrostenedione (11OHAD), to 11β-hydroxy-epi-testosterone (de Prada et al. 1994). Thus, important phenotypic differences exist relating to steroid metabolism between strains of *C. scindens*. de Prada et al. (1994) partially purified the native 17ɑ-HSDH from androstenedione-induced cultures of *C. scindens* VPI 12708 through ion exchange followed by two rounds of cibacron blue affinity chromatography and sequenced the N-terminus (de Prada et al. 1994). Almost thirty years later, we performed transcriptomic analysis of androstenedione-induced cultures of *C. scindens* VPI 12708 and identified a single gene that was significantly induced (GGADHKLB_RS03875; 3.07 log_2_FC; FDR 0.0099) (Wang et al. 2024). We cloned GGADHKLB_RS03875, overexpressed and affinity purified the recombinant protein and determined that this enzyme converts AD to epiT in an NADPH-dependent manner (**Figure 8**) (Wang et al. 2024). We named this gene the *desF* gene. Phylogenetic analysis indicated that *desF* is unique to *C. scindens* strains. Phylogenomic analysis indicated that *C. scindens* strains form separate clusters (12708 cluster and 35704 cluster), and average nucleotide identity (ANI) analysis suggests that *C. scindens* VPI 12708 and *C. scindens* ATCC 35704 may represent separate species (Caicedo et al. 2024). All strains harboring *desF* were in the 12708 cluster, and two strains encoded *desABC* and *desF* genes (Wang et al. 2024).

We then wanted to determine potential effects of epitestosterone formation on host physiology and health. Epitestosterone has long been regarded as an “antiandrogen”, a compound that antagonizes the nuclear androgen receptor (AR) (Maucher et al. 1994). A recent report indicates that the 5ɑ-reduced form of epitestosterone is an AR agonist in a reporter cell line (Schiffer et al. 2024). We reported that epitestosterone causes prolonged AR-dependent proliferation of prostate cancer cell lines that harbor a mutant AR (LNCaP) and wild type AR (VCaP) (Wang et al. 2024). In addition, the steroid-17,20-desmolase pathway is capable of side-chain cleave prednisone, used to treat prostate cancer, to 1,4-androstenedione (AT) as well as the 17ɑ-reduced form, epiAT (Wang et al. 2024). Addition of epiAT formed in spent medium by *C. scindens* VPI 12708 is able to promote significant proliferation of LNCaP cells. Furthermore, the drug abiraterone acetate (prescribed along with prednisone) is used to block adrenal androgen formation through the inhibition of host steroid-17,20-desmolase (CYP17A1) (Petrunak et al. 2023). Our results indicate that neither abiraterone acetate nor abiraterone are able to inhibit bacterial steroid-17,20-desmolase (Wang et al. 2024). We also measured fecal *desF* in patients currently on abiraterone acetate/prednisone that were responding (blood PSA levels stable) and when they became non-responsive (blood PSA levels rising) indicating androgens were increasing in tumors. We identified a subset of patients with fecal *desF* levels that increased when they went from responding to non-responding. A recent study also observed cortisol metabolism by *C. scindens* ATCC 35704 resulted in androgen-dependent LNCaP proliferation (Bui et al. 2023). Overall, these results indicate that *C. scindens* strains harboring *des* pathway genes may be important in prostate cancer progression, and potentially play a role in other androgen-dependent physiological and pathophysiological processes.

## *C. scindens* and the in vivo environment

While our knowledge of the pangenome of *C. scindens* is moving forward rapidly, our understanding of the microbial ecology of *C. scindens* is woefully behind. It is now clear that *C. scindens* is among a handful of bacterial species with “high” conversion rates of CA to DCA and CDCA to LCA (Ridlon et al. 2023). That the *bai* operon, encoding enzymes in the Hylemon-Björkhem Pathway, is part of the core genome of *C. scindens* affirms the importance of bile acid metabolism for this resident bacterial species in the human gut. Yet, the concentrations of *C. scindens* are relatively low in the large intestine of healthy humans (10^3^ to 10^5^ g^-1^ wet weight), while a few logs higher in gallstone patients (10^5^ to 10^7^ g^-1^ wet weight) (Berr et al. 1996). Nonetheless, even in low abundance, its potential to influence the health and well-being of the host cannot be underestimated. Recent evidence indicates that host-range goes beyond humans, with *C. scindens* strains isolated from both rat (Song et al. 2021) and pig (Wylensek et al. 2020).

Recent studies have examined the biogeographical distribution of *C. scindens* ATCC 35705 in gnotobiotic mice colonized with the OligoMM^12^, which harbors bacteria with bile salt hydrolase activity capable of forming free primary bile acids but lacks a member capable of bile acid 7ɑ-dehydroxylation (Marion et al. 2020). Previous in vitro studies established that *C. scindens* ATCC 35704 is not capable of bile salt hydrolysis and requires a free C24 carboxyl group in order to carry out bile acid 7ɑ-dehydroxylation (White et al. 1980). Interestingly, recent work suggests that *C. scindens* strains are capable of conjugating amino acids to bile acids, although the enzyme responsible for this is not clear (Guzior et al. 2024). Thus, in animals that are mono-colonized with *C. scindens* or in defined consortia lacking bile salt hydrolase activity (Narushima et al. 1999), the fecal bile acid profile would match the germ-free condition consisting of primary bile acids conjugated with taurine. Also, unlike rodent gut microbiota, human gut bacteria have not been shown to be capable of 7ɑ/7β-dehydroxylating MCA (Sacquet et al. 1984, Ridlon et al. 2020); whereas rat feces isolates convert murideoxycholic acid (MDCA) to MCA (Eyssen et al. 1999).

Marion et al. (2019) used nanoscale secondary ion mass spectrometry (NanoSIMS) to quantify *C. scindens* cultures before oral gavage in medium with ^15^N-labeled nutrients. Isotopically labeled *C. scindens* cells were detected 9 h after gavage in the distal intestine. *C. scindens* was present at 10^2^-10^3^ CFU g^-1^ in the ileum and 10^4^-10^7^ CFU g^-1^ in the cecum and colon at 24 h. These results are consistent with previous studies that report bile acid 7ɑ-dehydroxylating activity in the range of 10^3^-10^7^ per g^-1^ wet weight human stool (Berr et al. 1996). In vivo colonization with OligoMM^12^ and *C. scindens* resulted in a bile acid profile consistent with prior studies of germ-free mice “humanized” with patient stool (Berr et al. 1996) as well as the B3PC2 consortium which contains the bile acid 7α-dehydroxylating bacteria *C. hylemonae* and *C. hiranonis* (Narushima et al. 2006, Ridlon et al. 2020, Wolf et al. 2021). Taurine-conjugated primary bile acids were deconjugated in the cecum; however, only CA was converted to the secondary product DCA. Murine primary bile acids were not converted to MDCA (Marion et al. 2019).

In a follow-up study, Marion (2020) sought to determine the longitudinal distribution of bile salt biotransformation in the OligoMM^12^ with and without *C. scindens* ATCC 35704 (Marion et al. 2020). Metaproteomics and bile acid metabolomics was applied to each intestinal compartment demonstrating that addition of *C. scindens* to OligoMM^12^ affected species distribution and bile salt metabolism along the small and large intestines. *C. scindens* colonization in the OligoMM^12^ consortium led to decreased bile salt deconjugation in ileum, less bile salt hydrolase in proteome, and increased tauro-β-muricholic acid (TβMCA):β-MCA ratio. Low levels of tauro-DCA (TDCA) and tauro-MDCA (TMDCA) were detected in the liver, jejunum, and ileum only in mice colonized with *C. scindens* ATCC 35704. In the cecum and colon, *C. scindens* colonized mice exhibited DCA, LCA, MDCA, 12-oxo-LCA, and 6-oxo-alloLCA; whereas in the absence of *C. scindens* only oxo-primary bile acids were detected owing to HSDH enzymes expressed by OligoMM^12^ consortium members. These studies establish colonization biogeography with *C. scindens* ATCC 35704 and verify prior estimates of abundance and secondary bile acid production in vivo. Future studies are needed to understand the biology of *C. scindens* in the context of host-microbe and microbe-microbe interactions that determine colonization and abundance, and how these change with bile salt concentrations and dietary composition.

*C. scindens* ATCC 35704 has been shown recently to be capable of side-chain cleavage of glucocorticoid drugs such as dexamethasone and prednisone (Zimmermann et al. 2019, Ly et al. 2020). The side-chain cleavage product of prednisone was shown to stimulate growth of prostate cancer cells significantly greater than the most potent endogenous androgen, dihydrotestosterone (DHT) (Ly et al. 2020). Zimmermann et al. (2019) demonstrated steroid-17,20-desmolase activity against dexamethasone both in vitro and in vivo. Monocolonization of GF mice with *C. scindens* ATCC 35704 resulted in 10^9^ CFU g^-1^ content in the colon. Side-chain cleavage product of dexamethasone was observed to accumulate significantly in cecum and serum of *C. scindens* ATCC 35704 mono-associated mice relative to control mice (Zimmermann et al. 2019). 11-oxy-androgens, such as those generated by *C. scindens* ATCC 35704, have been a topic of increasing interest in the endocrine field due to their potential to signal through the androgen receptor (Bloem et al. 2013, Swart et al. 2015). The importance of strains of *C. scindens* that express steroid-17,20-desmolase on host physiology has yet to be explored.

## Interactions between *C. scindens* and *C. difficile*

Despite its relatively low abundance in the gut microbiome, *C. scindens* is likely to exert an inordinate role in maintaining microbiome structure through secondary bile acid production, and prevention of opportunistic pathogen colonization. Antibiotic-associated diarrhea caused by *C. difficile* is a growing global health threat with >450,000 infections and 29,000 death per year at a cost of roughly $5 billion in the U.S. alone (Lessa et al. 2015). Hospitals are a major source of infection due to higher environmental load of *C. difficile* spores coupled with a population having a greater probability of antibiotic use. *C. difficile* spores germinate in the gastrointestinal (GI) tract producing toxin A and B from secreting vegetative cells that cause symptoms ranging from diarrhea to severe colitis (Schnizlein et al. 2022). Metronidazole or vancomycin are used to initially treat *C. difficile* infection. However, roughly 10-40% of patients that are successfully treated relapse following the end of antimicrobial therapy (Schnizlein et al. 2022). Fecal microbiota transplants (FMT) from healthy donors have been shown to be highly effective in treating patients relapsing from *C. difficile* treatment, indicating reestablishment of normal microbial inhabitants is necessary to exclude *C. difficile* from the GI environment. Research efforts in recent years have focused on determining which gut microbial species are both necessary and sufficient to treat *C. difficile* infection, providing targeted, defined probiotic alternatives to FMT (Lavoie et al. 2023).

Bile acids are thought to be central to *C. difficile* germination (**Figure 9**). Indeed, *C. difficile* spores require 12ɑ-hydroxylated bile acids, principally taurocholic acid (TCA) to germinate in vitro (Sorg et al. 2008, Francis et al. 2013). TCA in the presence of glycine, released during microbial bile salt hydrolysis, enhances *C. difficile* germination (Sorg et al. 2008). Francis et al. (2013) demonstrated that the subtilisin-like pseudoprotease, CspC, is the germination receptor that recognizes 12ɑ-hydroxylated bile acids. In contrast, bile acids that lack a 12ɑ-hydroxyl group (e.g., CDCA and MCA) prevent *C. difficile* spore germination (Sorg et al. 2010) and act as competitive inhibitors of TCA-induced germination (Sorg et al. 2010, Francis et al. 2013). Binding of bile acids is predicted to exchange Ca^2+^-dipicolinic acid for water in the spore cortex, allowing for cell wall hydrolysis and vegetative growth.

**Figure 9.**
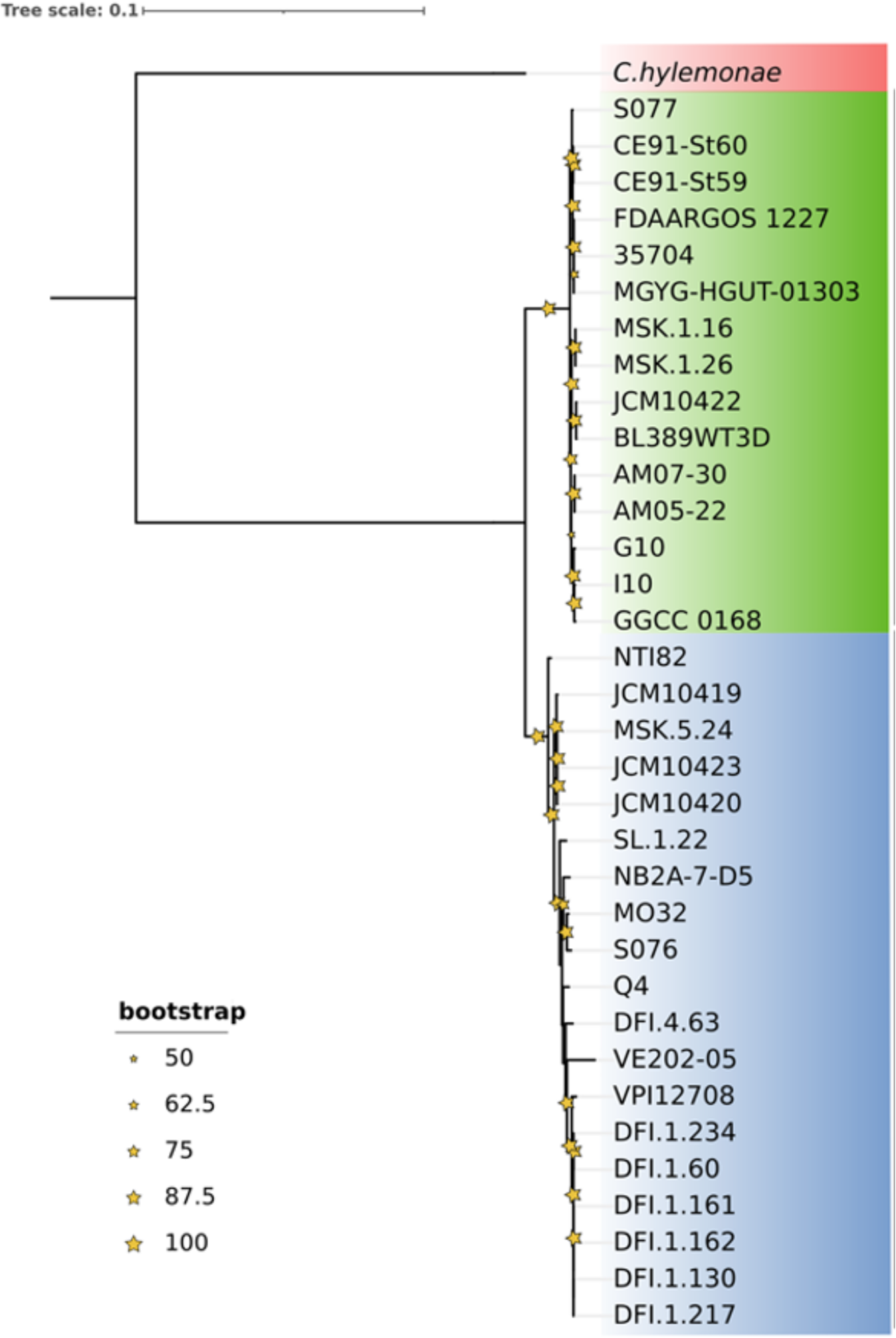
The role of bile acid and tryptophan metabolism in germination and vegetative growth of *Clostridioides difficile*. Taurocholic acid is deconjugated, mainly in the large intestine, by diverse gut microbial taxa. Free cholic acid is imported into a few species of Bacillota that harbor the *bai* regulon. Direct Pathway: After several oxidative steps, and rate-limiting 7α-dehydration, 3-oxo-Δ^4^-DCA becomes a substrate for BaiCD forming DCA or BaiP/BaiJ forming alloDCA. Indirect Pathway: DCA is imported into Bacteroidetes strains that express 3α-HSDH and 5β-reductase (5BR) which converts DCA to 3-oxo-Δ^4^-DCA. Expression of 5α-reductase (5AR) and 3β-HSDH sequentially reduce 3-oxo-Δ^4^-DCA to iso-allo-DCA. Allo-DCA generated by Bacillota is also isomerized to iso-allo-DCA via 3α-HSDH and 3β-HSDH expressing strains of *Eggerthella lenta* and other taxa. While TCA is a germination factor for *C. difficile,* DCA and isoalloLCA have been shown to be inhibitory toward *C. difficile* vegetative growth in vitro and in vivo. Secondary bile acids, including DCA and allo-DCA, are associated with increased risk of colorectal cancer (CRC). In addition, *C. scindens* strains have been shown to convert L-tryptophan to 1-acetyl-β-carboline which promotes synergistic inhibition of *C. difficile* in the presence of hydrophobic secondary bile acids (). Modified from a previously published image with kind permission ().

Bile acids have also been shown to affect *C. difficile* growth in vitro (**Figure 9**). Secondary bile acids such as DCA and LCA have been reported to impair vegetative cell growth in vitro (Wilson 1983, Sorg et al. 2008). Theriot et al. (2016) demonstrated in *ex vivo* ileal and cecal contents from mice treated with cefoperazone allowed *C. difficile* spore germination while conventional content prevented germination. Intestinal content from cefoperazone treated mice were significantly depleted in unconjugated secondary bile acids.

In 2015, *C. scindens* was identified as top among a handful of microbial taxa associated with resistance to *C. difficile* infection in mice and humans (Buffie et al. 2015). Adoptive transfer of a four-strain consortium of microbes predicted to inhibit *C. difficile* which included *C. scindens* or *C. scindens* alone, was capable of ameliorating *C. difficile* infection in a murine model treated with antibiotics (Buffie et al. 2015). Indeed, recovery of antibiotic-treated mice from *C. difficile* positively correlated with recovery of secondary bile acids and the *baiCD* gene. Bile acid-binding anion-exchange resin cholestyramine treatment permitted *C. difficile* growth, which was interpreted as demonstrating bile acid-dependent inhibition. Subsequent clinical studies supported this association, observing a negative association between fecal *baiCD* abundance and *C. difficile* infection (Solbach et al. 2018). Importantly, secondary bile acids have varying degrees of growth inhibition against *C. difficile* in vitro (Giel et al. 2010, Kang et al. 2019, Sato et al. 2021). In particular, derivatives of planar bile acids, particularly isoallolithocholic acid appear to be particularly inhibitor even at low micromolar concentrations (**Figure 9**) (Sato et al. 2021).

In addition to the formation of growth-inhibitory secondary bile acids, *C. scindens* ATCC 35704 also synthesizes the antimicrobial compound, 1-acetyl-beta-carboline which inhibits cell division of *C. difficile* (Kang et al. 2019). Likewise, *C. difficile* ATCC 9689 and clinical isolates (BBA-1870, BBA-1801, and BBA-1814) produce cyclo(Phe-Pro) and cyclo(Leu-Pro) dipeptides that inhibit the growth of *C. scindens* ATCC 35704 (Kang et al. 2019). During *C. difficile* infection, host collagen is degraded by metalloproteases in response to CDI toxins, and this results in release of posttranslationally modified trans-4-hydroxyproline (Reed et al. 2022). *C. difficile* competes with *C. scindens* VPI 12708 in vivo for proline (Reed et al. 2022). Indeed, Cyp8b1-/- (cholic acid-deficient) mutant mice are protected from infection by *C. difficile* spores in the presence of *C. scindens* VPI 12708 in mono-associated gnotobiotic mice (Aguirre et al. 2021). The production of 1-acetyl-beta-carboline was not detected in gnotobiotic mouse or patient fecal samples, and metabolomics analysis suggests that competition in vivo for the Stickland fermentation of proline is important, rather than bile acid metabolism (Aguirre et al. 2021). *C. difficile* also shares several nutritional requirements with *C. scindens* such as the amino acid tryptophan and the B vitamins pyridoxine and pantothenate (Karasawa et al. 1995, Devendran et al. 2019). These results suggest that some combination of bile acid metabolites, antibiotic warfare, and competition for nutrients determines the success of *C. difficile* infection vs. gut microbial homeostasis.

## The pangenome of *C. scindens*

The past 40 years of research on *C. scindens* has been relegated to the two strains reported in the early 1980s, *C. scindens* VPI 12708 and *C. scindens* ATCC 35704. However, the importance of strain variation among members of the human microbiome cannot be underscored enough (Britton et al. 2021). A partial genome for *C. scindens* ATCC 35704 was reported as part of the Human Microbiome Project (PRJNA18175) in 2006. Recently, we reported the complete 3,658,040 bp genome of *C. scindens* ATCC 35704 which comprised 3,657 coding sequences (CDS), 12 rRNA genes, 4 rRNA cistrons, and 58 tRNA genes (Devendran et al. 2019). Annotation of the genome indicated certain nutritional requirements due to the absence of genes involved in the de novo synthesis of tryptophan, riboflavin, pyridoxal phosphate, and panthothenic acid (Devendran et al. 2019). The partial genome of *Clostridiales* VE202-05 (PRJDB524) appears to be closest to the genome of *C. scindens* VPI 12708, which we recently sequenced (Caicedo et al. 2023). *C. scindens* ATCC 35705 and VPI 12708 share ∼64.5% of their genes (Devendran et al. 2019). This is in contrast to our recently reported closed genome of *C. scindens* BL389WT3D isolated from swine feces which shared 81.9% of their 3,656 CDS with the type strain ATCC 35704 (Wylensek et al. 2020). One of the interesting findings from comparing the genomes of the human ATCC 35704 and pig BL389WT3D genomes is that of the ∼660-690 unique genes, much of this content is composed of mobile genetic elements such as bacteriophage genes and transposons (Wylensek et al. 2020).

We recently analyzed the genomes of 34 cultured strains of *C. scindens* (**Table 3**). These include 9 additional sequenced genomes that we recently reported (Fernandez-Materan et al. 2024), 8 complete genomes obtained from the public GenBank database at the National Center for Biotechnology Information (NCBI), and 17 incomplete genomes at the level of contigs and scaffolds obtained from NCBI. In addition, the genomes of 66 assembled metagenomic genomes (MAGs) of *C. scindens* from the intestinal metagenome of human fecal samples were included (Pasolli et al. 2019, Almeida et al. 2021, Zeng et al. 2022). The analysis identified a pangenome with 12,720 gene families, distributed in three groups, and associated with the core genome, accessory genome, and unique or strain-exclusive genes. A total of 1,630 gene groups are in the core, representing almost 13% of the total pangenome, 7,051 accessory groups, and 4,039 unique genes (Caicedo et al. 2024). On the other hand, the accessory genome was determined using MAGs genomes with a completeness value equal to or greater than 85%. A total of 157 genomes shows a pangenome size of 19,198 gene families and a core genome of 132 gene groups, the core representing almost 7% of the total pangenome. The application of Heap’s Law formula demonstrated that the pangenome was open when the 34 cultured strains were included (ɑ = 0.845) and remained open when 66 MAGs of *C. scindens* were included (ɑ = 0.768). Phylogenomic analysis of the 34 strains revealed two clusters: a 12708 group and a 35704 group (**Figure 10**).

**Figure 10.**
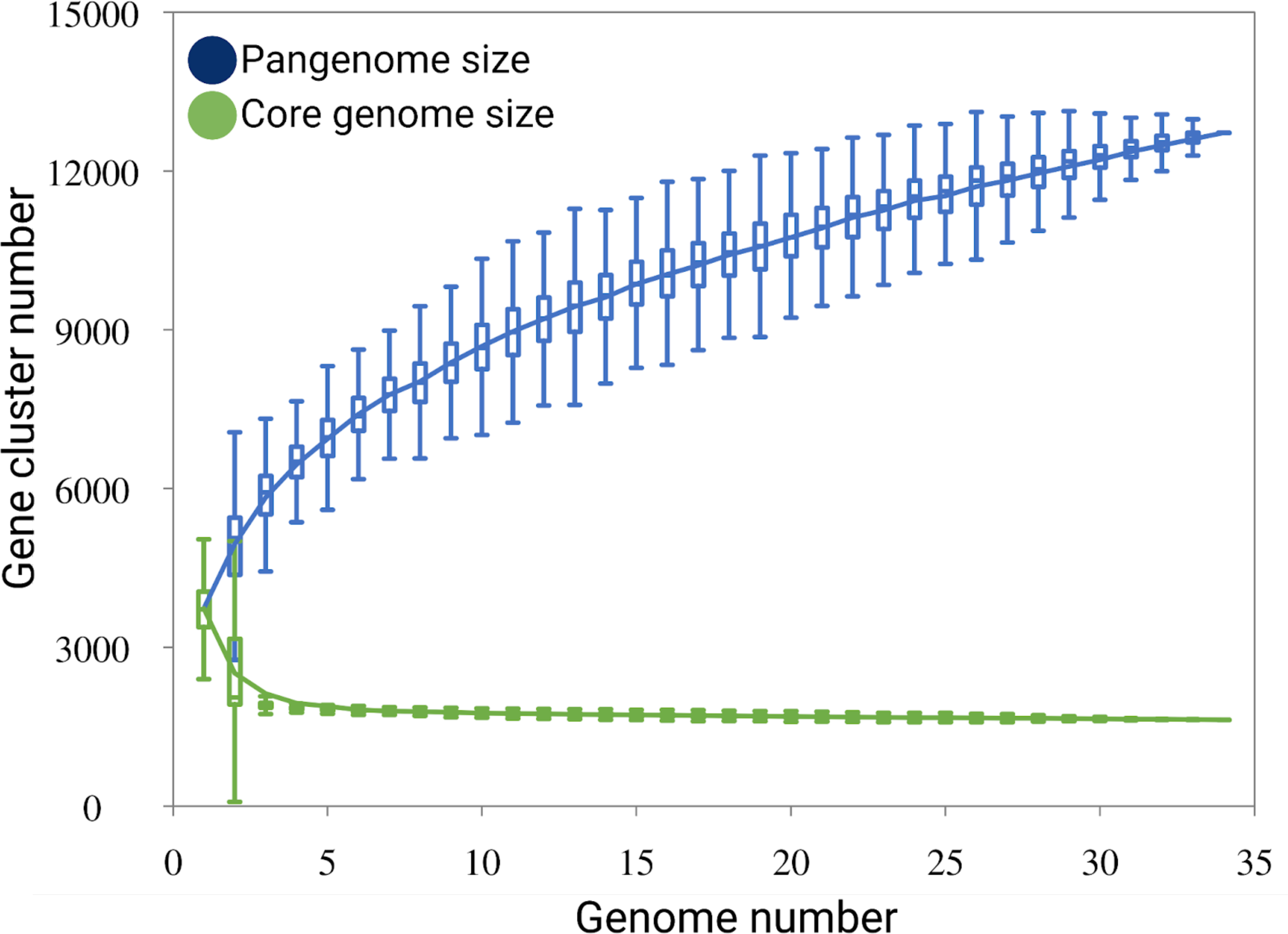
Phylogenomic and diversity of *des* gene presence in strains of *C. scindens*. The formation of two clades is shown, Clade 1 (green) includes 15 strains and Clade 2 (blue) 19 strains. Bootstrap support values above 50% are shown in yellow stars at nodes.

**Table 3.**
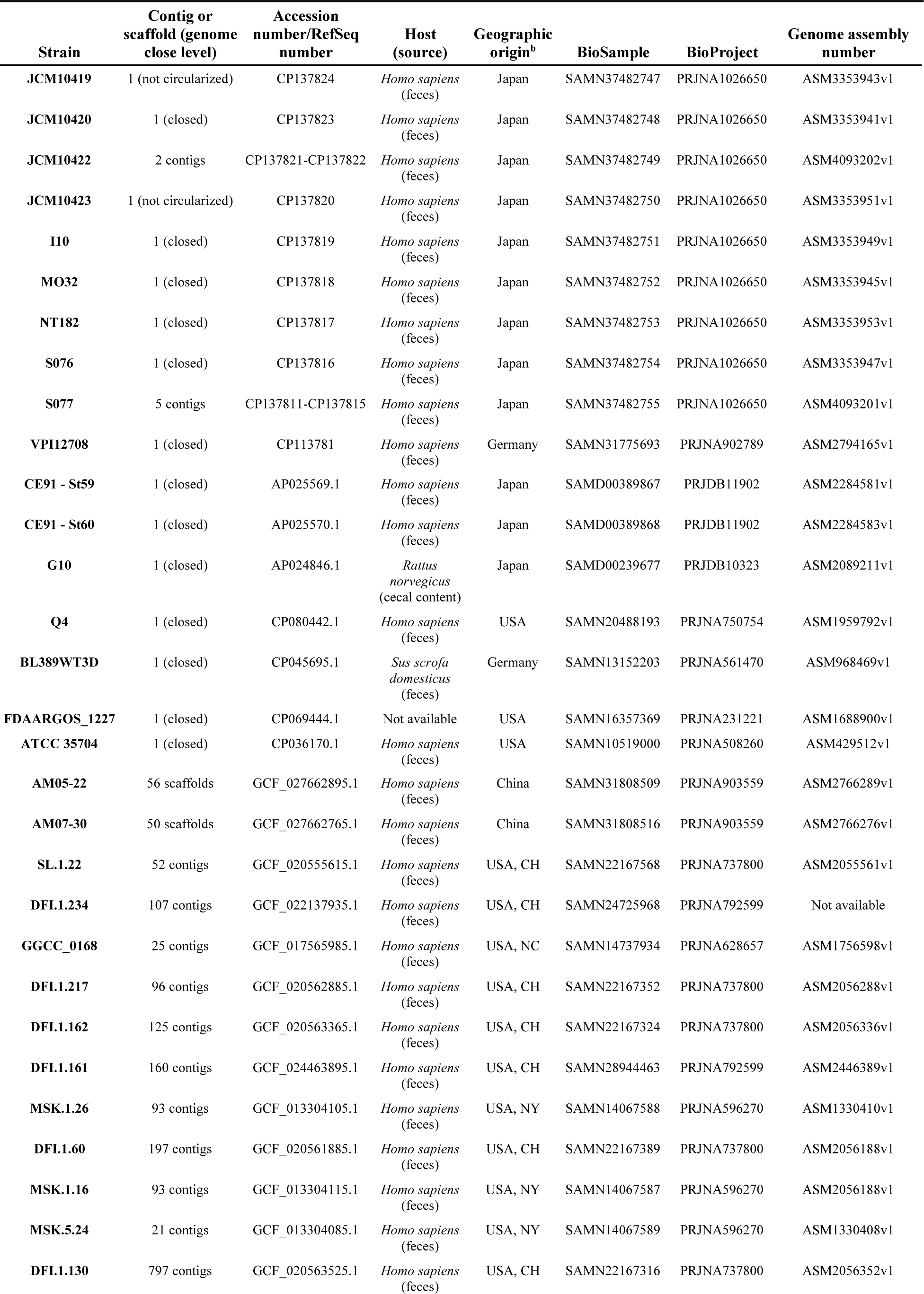
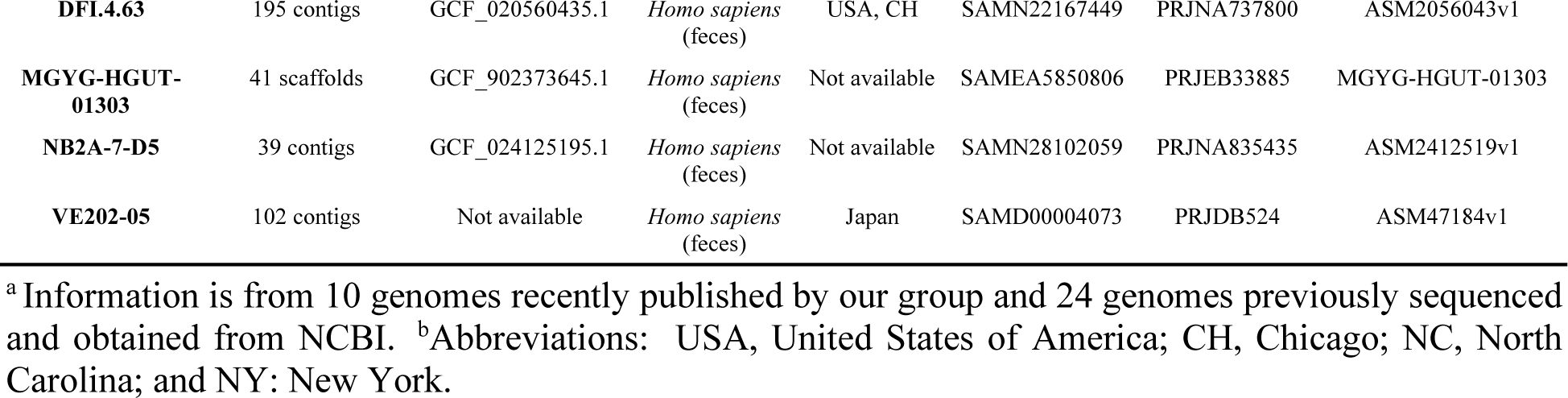
Characteristics and genomic information for 34 cultured strains of *C. scindens*.^a^.

Average nucleotide identity (ANI) analysis between the two strain groups was then performed. The suggested threshold percentage value for ANI analysis of species identity is equal to or greater than 95% (Richter et al. 2009), identified two sets of strains. The intraspecies delineation criterion was also considered through the analysis of distances between genomes. The results show both isolated groups divided into 15 and 19 strains with a difference of approximately 4 to 5% in their genomic sequences. Identity within each group is ≥ 98%, while identity between groups is 94.5 to 96%, whereas with *C. hylemonae* genome the identity values were between 74 to 76%. The present ANI values of *C. scindens* suggest the presence of two possible distinct bacterial species or at least an ongoing speciation process. Generally, in a set of strains belonging to the same species, the spectral function of the pangenome is “U’’ shaped without internal peaks that differentiate the number of gene groups found in the genomes, but in a small number of species mixed sample, the spectrum function will have internal peaks (**Figure 11**). In other words, it is defined as “homogeneous”, a set of genomes with a “U” shaped spectrum function, and “non-homogeneous”, a the set with internal peaks (Moldovan et al. 2018). Consequently, the gene group frequency plot shows the presence of internal peaks in the accessory genome, corresponding to a non-homogeneous group of strains, possibly produced by the gain and/or loss of genes or by the fusion of two homogeneous groups, where the central peaks of the G(k) function correspond to specific genes for homogeneous groups.

**Figure 11.**
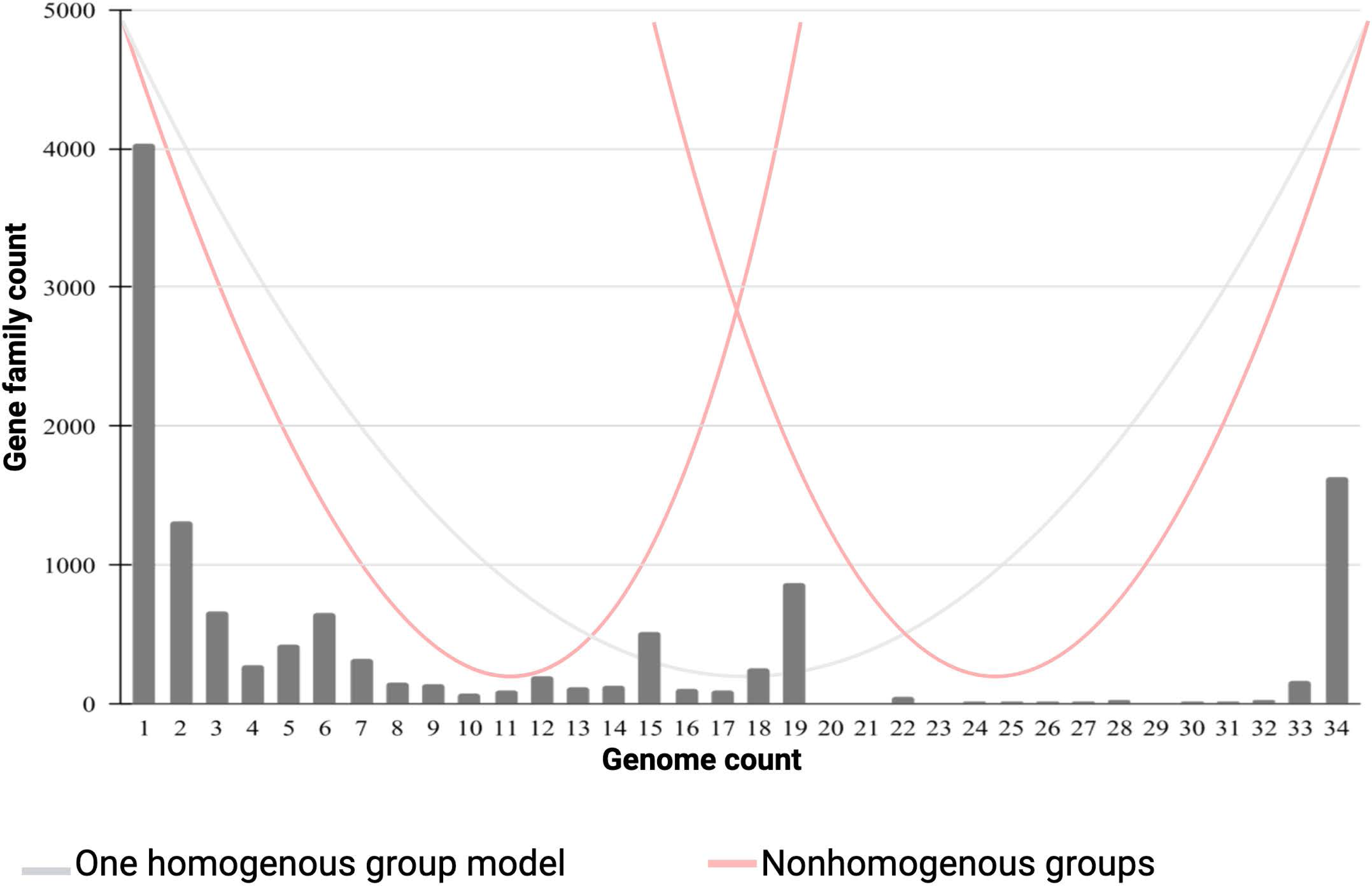
Gene frequency versus number of *C. scindens* genomes. **A.** Pan-core plots for *C. scindens.* **A.** Pan-core plot of the 34 cultured strains of *C. scindens*. The graphics show cumulative curves of the upward trend in the number of pangenome gene families (in blue) and the downward trend of core gene families (in green) with each consecutive addition of a *C. scindens* genome. The rising curve in blue shows an open pangenome. **B.** The graph shows the frequency of gene clusters in the 34 *C. scindens* genomes, the left bar “1” represents the number of strain-specific genes, the right bar “34” indicates the core genome, the central bars refer to the accessory genome. Lines are drawn to show the theoretical distribution of a “U” shaped “homogeneous” distribution (gray) and a “W” shaped distribution (red) for “non-homogeneous” plot.

## Predicted metabolic pathways in the core genome

The complete Hylemon-Björkhem pathway is a core feature of *C. scindens* strains (**Figure 12**). While steroid-17,20-desmolase (*desAB* genes) is present in both groups, it is by far more prevalent in Group 1 (35704 group) (**Figure 13**). By contrast, Group 2 (12708 group) had sole representation of the *desF* gene, including two strains that have both *desABCD* and *desF* genes (**Figure 13)**. The 12708 group also has sole representation of the *baiJ* and *baiK* genes, previously shown to encode bile acid 5ɑ-reductase (Lee et al. 2022) and bile acid CoA transferase (Ridlon et al. 2012), respectively. Overall, these findings tend to support Bokkenheuser’s claims of “taxonomic value” for steroid-metabolizing activities (Bokkenheuser 1993), thereby suggesting that *C. scindens* VPI 12708 and ATCC 35704 may very well represent distinct and separate species. What the taxonomic designation should be for these two groups (clades) awaits determination.

**Figure 12.**
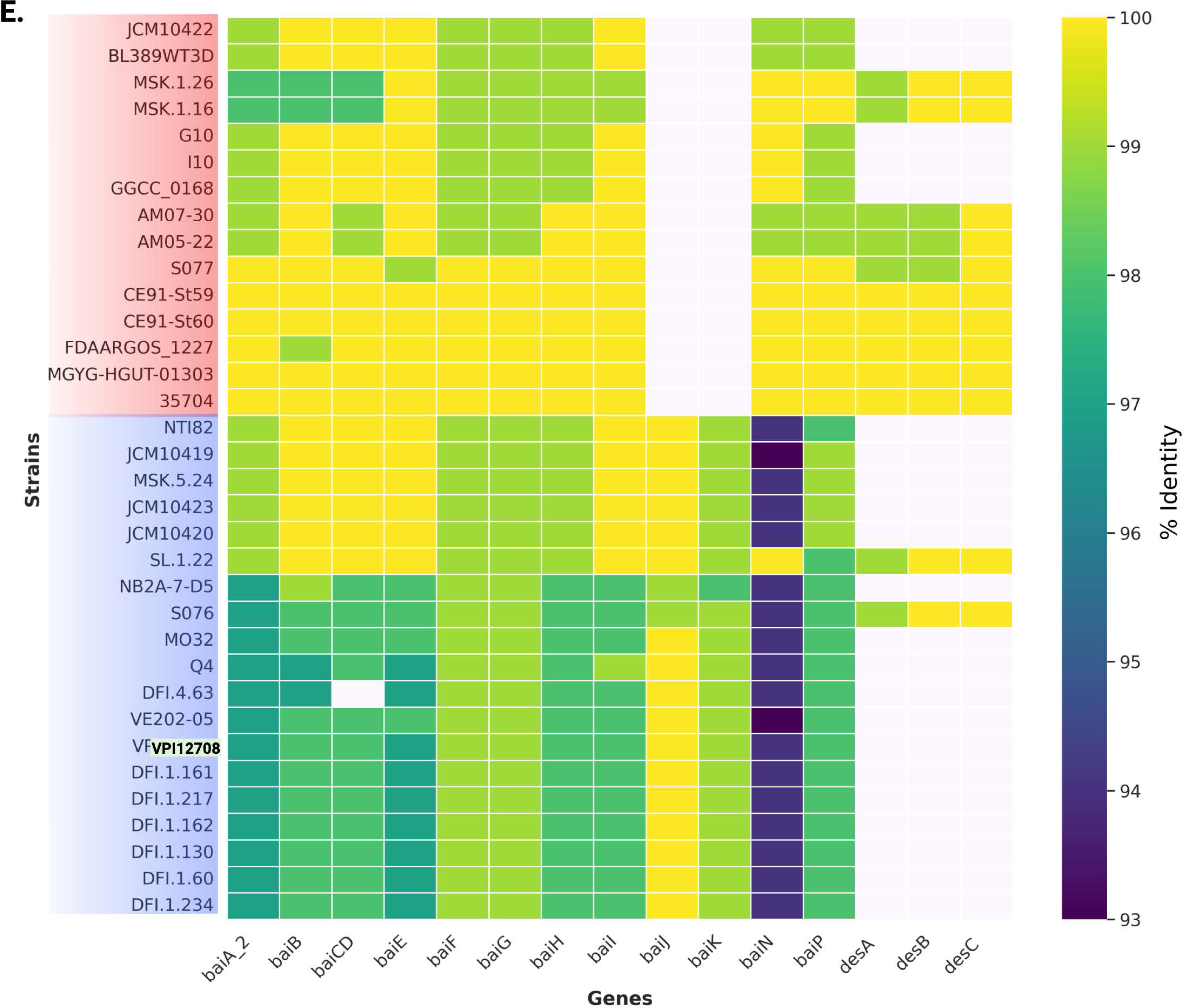
Heatmap of *bai* and *des* genes in the 34 *Clostridium scindens* genomes. The percentage value of sequence identity is shown in color; the highest value in yellow and the lowest in purple. The lack of the gene is indicated by a white rectangle.

**Figure 13.**
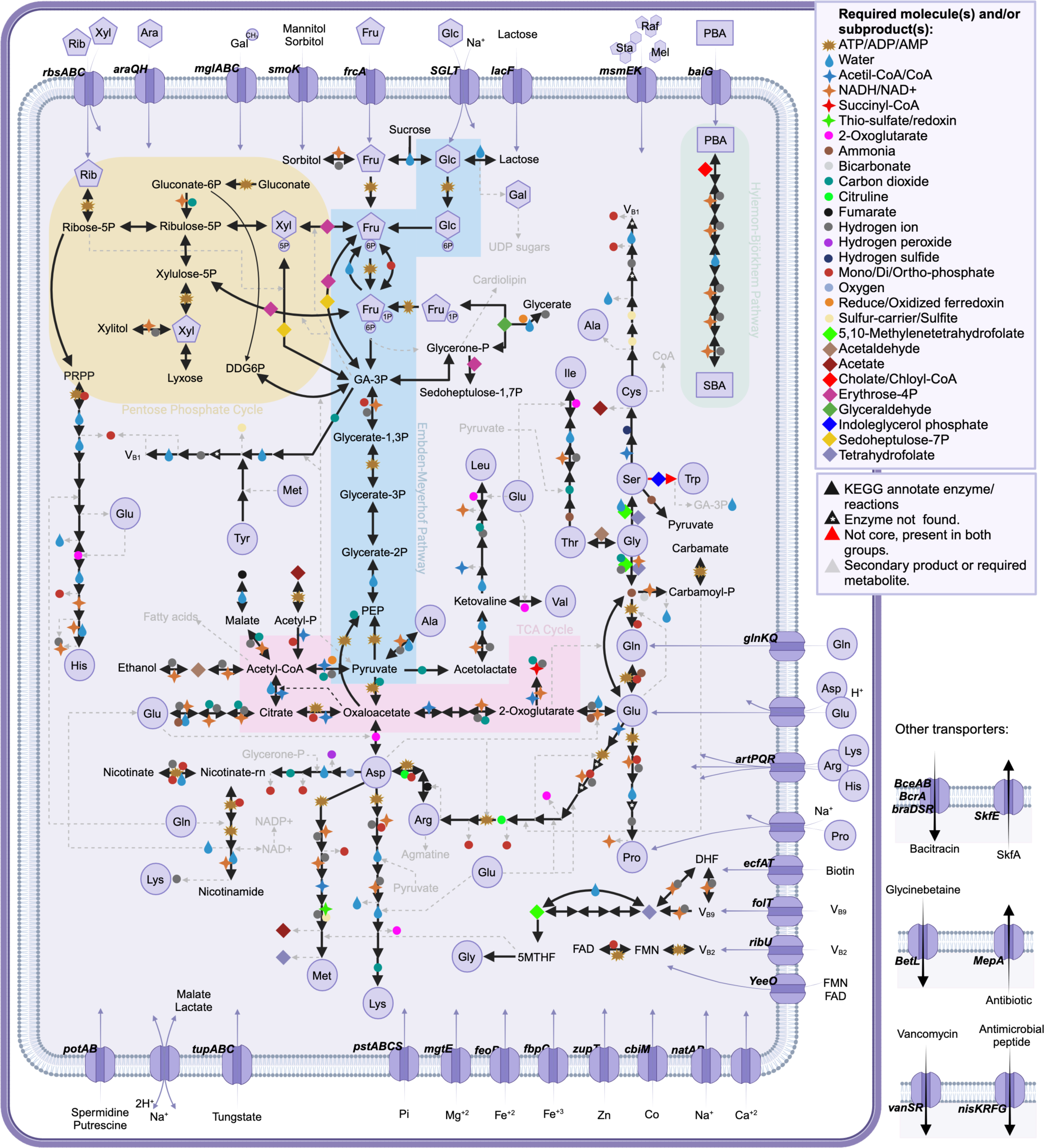
Key metabolic pathways represented in the core genome of *Clostridium scindens*.

Overall, *C. scindens* strains harbor the complete glycolytic pathway as well as Entner-Doudoroff Pathway. A complete pentose phosphate pathway, and a “horseshoe” TCA cycle from oxaloacetate to succinyl∼SCoA are also observed where oxaloacetate is generated from phosphoenolpyruvate, malate and fumarate from pyruvate. The core genome contains complete pathways for the biosynthesis of the majority of amino acids. The core and accessory genomes contain the near complete shikimate pathway for the biosynthesis of L-phenylalanine and L-tyrosine; however, an enzymatic pathway to tryptophan is not observed, nor is a *de novo* pathway for the synthesis of proline found in the core genome.

While a pathway from pantothenate to CoA was evident in the core genome, pantothenate biosynthesis is not present. (**Figure 8**). While de novo nicotinate biosynthesis is lacking, genes encoding enzymes from nicotinate to NAD+ and NADP+ are present. Thiamine and cobalamin biosynthesis pathways are part of the core genome, as are genes encoding enzymes involved in one carbon pool by folate. Genes encoding enzymes involved in converting riboflavin to FMN and FAD are present; however, biosynthesis pathways for riboflavin are absent. Lipoate salvage, but not biosynthesis, is present in the core genome. Thus, it is advised that our previous defined medium for *C. scindens* ATCC 35704 (Devendran et al. 2019) may require supplementation with a complete set of vitamins (except thiamine) and additional amino acids (tryptophan, proline, phenylalanine, tyrosine) in order to cultivate other *C. scindens* strains under defined conditions.

## Conclusions

*C. scindens* is a keystone gut microbial taxonomic group that, while low in abundance, has a disproportionate effect on bile acid and steroid metabolism in the GI tract. Both the Hylemon-Björkhem pathway and the steroid-17,20-desmolase pathway were first discovered in *C. scindens.* Numerous studies indicate that the two most studied strains of *C. scindens* (i.e., ATCC 35704 and 12708) are important for a myriad of physiological processes in the host. Our most recent analysis now calls into question whether strains currently defined as *C. scindens* represent two separate taxonomic groups. Future directions include developing genetic tools to further explore (i) the role of *bai* and *des* genes in steroid metabolism by *C. scindens*, (ii) the interaction between steroid metabolism and essential (core) metabolic pathways in *C. scindens* and its impact on carbon and reductant flow in this bacterium (**Figure 14**), and the causal role of steroid-metabolizing pathways by *C. scindens* in host physiology and disease.

**Figure 14.**
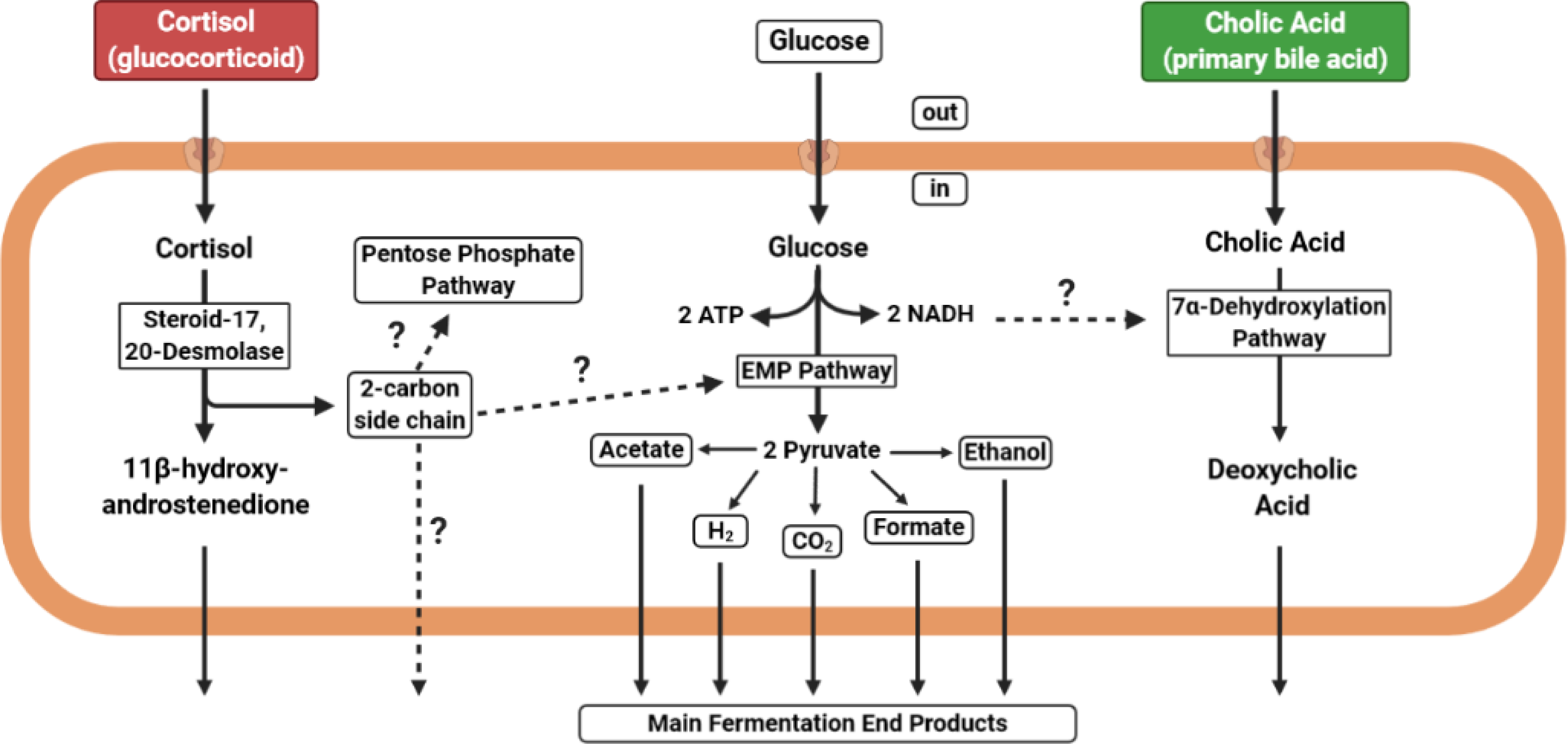
A proposed model for the interaction between glucose fermentation, cortisol metabolism, and bile acid 7α-dehydroxylation by *Clostridium scindens* ATCC 35704. EMP, Embden-Meyerhof-Parnas pathway.

## Acknowledgment

The authors would like to express their deepest appreciation to the following individuals: Carl Bokkenheuser for pictures and information about his uncle Victor D. Bokkenheuser; Michala Biondi, archivist at the Icahn School of Medicine at Mount Sinai in New York, for information about Victor D. Bokkenheuser during his time at St. Luke’s-Roosevelt Hospital Center in New York; Sheryl Locascio, Anne Mosenthal, Emilia Sordillo, and Irene Grant for information and their reflections on working with Victor Bokkenheuser; Jacqui Winter for the picture of her mother Jeanette Winter; Anna Cerone-McLernon for the picture of herself; George Morris for the picture of himself; Sheryl O’Rourke-Locascio for the picture of herself; Doris Hammann for the picture of her late husband Rainer Hammann; John Jackson at Special Collections and University Archives, University Libraries, Virginia Polytechnic Institute and State University for the pictures of Lillian Moore and W.E.C. Moore; Jeff Karr, archivist at the American Society for Microbiology, for the photo and information about Elizabeth Cato; Mark Kehrli, former Director of the National Animal Disease Center in Ames, Iowa, and Diana Whipple for photos of Alfred Ritchie; Steven Daniel for the picture of Phillip Hylemon and Bryan White; Amy Krafft for the picture of herself; Jen and Linda Macdonald for the picture of Ian Macdonald; Hebba Beech at the Microbiology Society for permission to reuse the electron micrograph photo of *Clostridium scindens* ATCC 35704; and Takeshi Katagiri from RIKEN and the Japanese Collection of Microorganisms for permission to reuse the Gram stain and colonies-on-plate photos of *Clostridium scindens* 35704. We would also like to thank Prof. Phillip B. Hylemon for helpful discussions relating to early collaborative efforts to characterize *C. scindens* strains. J.M.R. is supported by grants from the National Institute of Allergy and Infectious Disease (R03AI147127) and the National Institute of General Medicine (R01GM134423).

---

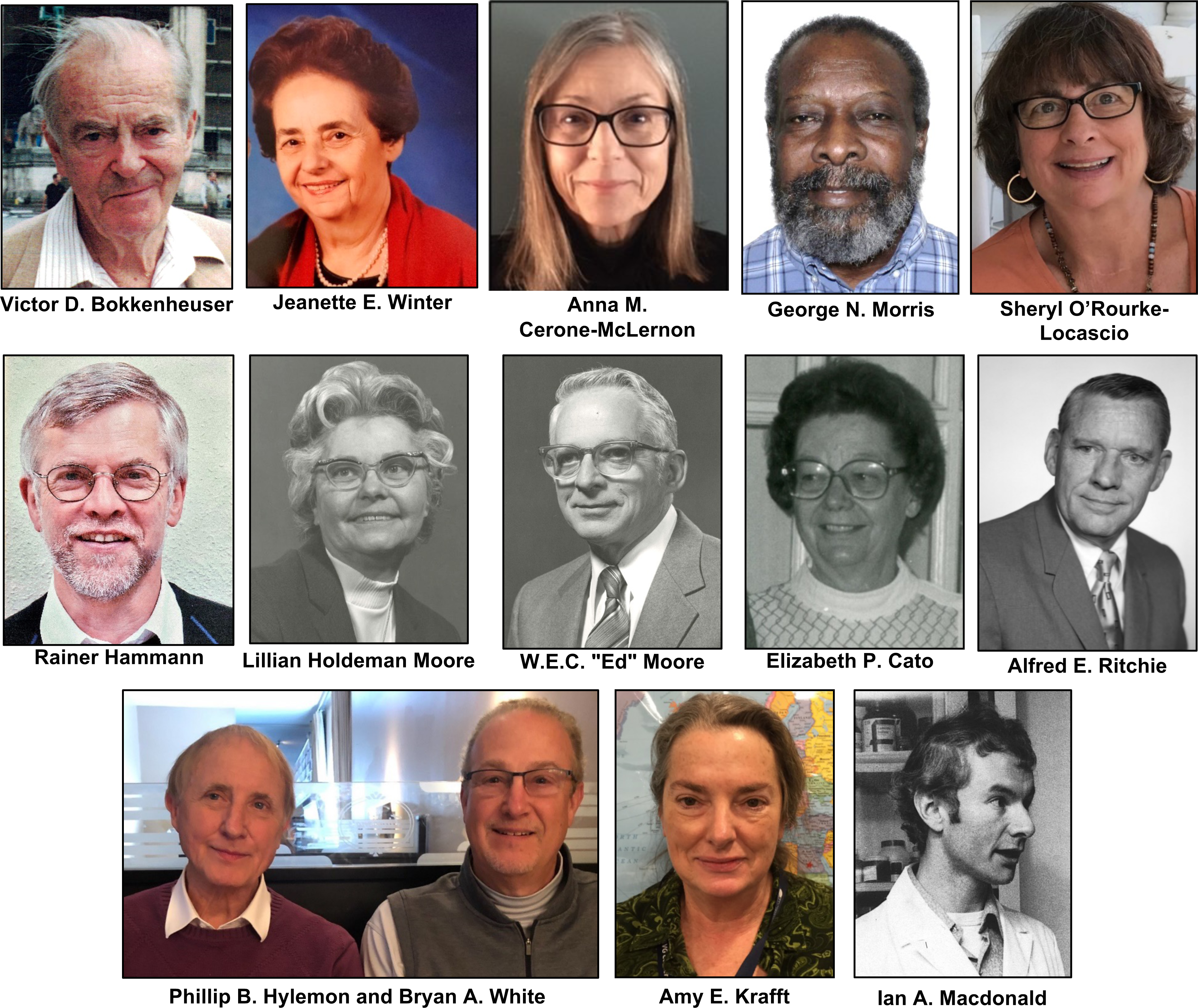

